# A membrane-targeted photoswitch restores physiological ON/OFF responses to light in the degenerate retina

**DOI:** 10.1101/2024.10.02.616298

**Authors:** Gaia Ziraldo, Sara Cupini, Valentina Sesti, Emanuela Delfino, Guglielmo Lanzani, Chiara Bertarelli, Fabio Benfenati, Stefano Di Marco

## Abstract

The lack of effective therapies for visual restoration in *Retinitis pigmentosa* and macular degeneration has led to the development of new strategies such as optogenetics and retinal prostheses. However, visual restoration is poor due to the massive light-evoked activation of retinal neurons, regardless of the segregation of visual information in ON and OFF channels, essential for contrast sensitivity and spatial resolution. Here, we show that Ziapin2, a membrane photoswitch which modulates neuronal capacitance and excitability in a light-dependent manner, is capable of reinstating, in two distinct genetic models of photoreceptor degeneration, brisk and sluggish ON, OFF, and ON-OFF responses in retinal ganglion cells evoked by full-field stimuli, with reactivation of their excitatory and inhibitory conductances. Intravitreally injected Ziapin2 in fully blind rd10 mice restored light-driven behavior and optomotor reflexes. The results indicate that Ziapin2 is a promising molecule for reinstating physiological visual responses in the late stages of retinal degeneration.

## INTRODUCTION

Vision relies on extracting and recognizing shapes from the environment. The ability of the retina to elaborate visual information is incomparably higher than that of the most advanced cameras available nowadays. This outstanding performance resides in the capacity of the retinal network to segregate visual information into distinct parallel channels to simultaneously process positive or negative contrast in the ON and OFF pathways, respectively, allowing to segregate and extract visual features, like spatial organization or movement. As a result, in the rodent retina, more than 30 functional types of retinal ganglion cells (RGCs) characterized by distinct responses to light and dark can be identified^1–3^. The segregation of visual information and contrast encoding are initially generated at the photoreceptor ribbon synapse by the inhibitory surround generated by horizontal cells and the physical segregation of positive/negative contrasts in three main classes of bipolar cells (BCs), with rod- and cone ON-BCs expressing hyperpolarizing metabotropic glutamate receptors 6 (mGluR6) and cone OFF-BCs expressing depolarizing AMPA/kainate glutamate receptors. Rod BCs of the peripheral retina modulate cone ON- and OFF-pathways in an opposite fashion, contributing to contrast sensitivity: they excite cone ON-BCs directly *via* gap junctions and inhibit cone OFF-BCs *via* AII-amacrine cells’ glycinergic synapses. From this point on, visual information remains segregated in these two parallel information streams until it reaches the higher integrative centers ^4–8^.

The progressive degeneration of retinal photoreceptors is one of the most frequent causes of severe visual impairment. *Retinitis pigmentosa* (RP), a collective name for a set of genetic disorders that cause the death of rods and secondarily of cones, afflicts 1 in 3,500/4,000 people worldwide and is the most common inherited retinal degeneration^9,10^. On the other hand, age-related macular degeneration (AMD), involving cones in the *fovea centralis* of the macula that are responsible for sharp central vision ^4^, affects ∼8% of the general world population and ∼25% of people above 70 years of age ^11^. Notwithstanding the prevalence of AMD and RP, therapeutic approaches have not yet been successful, and no approved pharmacological treatment exists ^12,13^.

In addition to gene and cell therapies aimed to counteract or slow down photoreceptor death ^14–23^, alternative strategies point to the light-dependent reactivation of the inner retinal networks spared by degeneration. Optogenetics, widely tested in preclinical models, was recently reported in one RP patient within an ongoing clinical trial ^24,25^. Despite the advantage of applying to any RP mutation and stage of the disease, the inherent low light sensitivity of heterologous opsins requires light intensification, and the resulting visual acuity is relatively scarce. Retinal prostheses generating light-induced electrical signals activating the inner retina and restoring some extent of light sensitivity in dystrophic retinas have been proposed ^26–35^. Within the eye, prosthetic chips have been implanted either in the epiretinal or subretinal position. Epiretinal devices, closer to RGCs, have the advantage of stimulating the final common pathway of the retina. At the same time, subretinal implants could make good use of the computational processing of the inner retina. However, not considering the intrinsic limitations of planar devices (tiny visual field and spatial resolution 1-2 orders of magnitude lower than macular cones), all prosthetic approaches, including those based on photovoltaic nanoparticles^36–38^, induce a light-evoked indiscriminate activation of the inner retina, compressing all visual information into the ON pathway, unable to generate negative contrast information and high spatial resolution.

A new generation of azobenzene-based photoswitches has recently been demonstrated to photosensitize endogenous proteins without the requirement of genetic engineering ^39,40^. The most recent photoswitches are sensitive to visible light and trigger light-reversible blockade of voltage- or ligand-gated ion channels of RGCs by acting intracellularly at the channel site ^41–48^. However, the inability to generate negative contrast information is also shared by these compounds that can only modulate the retinal network upon illumination, eliciting a massive activation of the ON pathway.

We have recently engineered and biologically characterized a novel amphiphilic azobenzene-based photoswitch, named Ziapin2, that acts on passive properties of the membrane, without directly interfering with ion channels or neurotransmitter receptors. Ziapin2 is composed of an azobenzene core flanked by two *pyridinum*-capped branches on one side and a hydrophobic azepane ring on the opposite side. Ziapin2 spontaneously partitions into both membrane leaflets, by aligning its polar termini with the phospholipid headgroups, and its hydrophobic core with the membrane hydrophobic lipid chains ^48^. In the dark, the two azepanes of opposite Ziapin2 molecules trans-interact and decrease membrane thickness, thus increasing capacitance and making neurons less excitable. The *trans-to-cis* isomerization triggered by millisecond pulses of cyan light (450-500 nm; peak at 470 nm; **Figure 1a**, left) causes membrane relaxation by removing the hydrophobic end-group from the membrane core, resulting in a sharp decrease in capacitance. This causes a fast hyperpolarization of the neuronal membrane, followed by a rebound depolarization. These changes increase intrinsic excitability due to an anode breaking mechanism and trigger action potential (AP) firing ^49–51^. The bidirectional dark/light modulation of neuronal excitability makes this molecule a promising tool for separating discharge areas from the inhibitory surround within the ON/OFF pathways of the retina.

**Figure 1.**
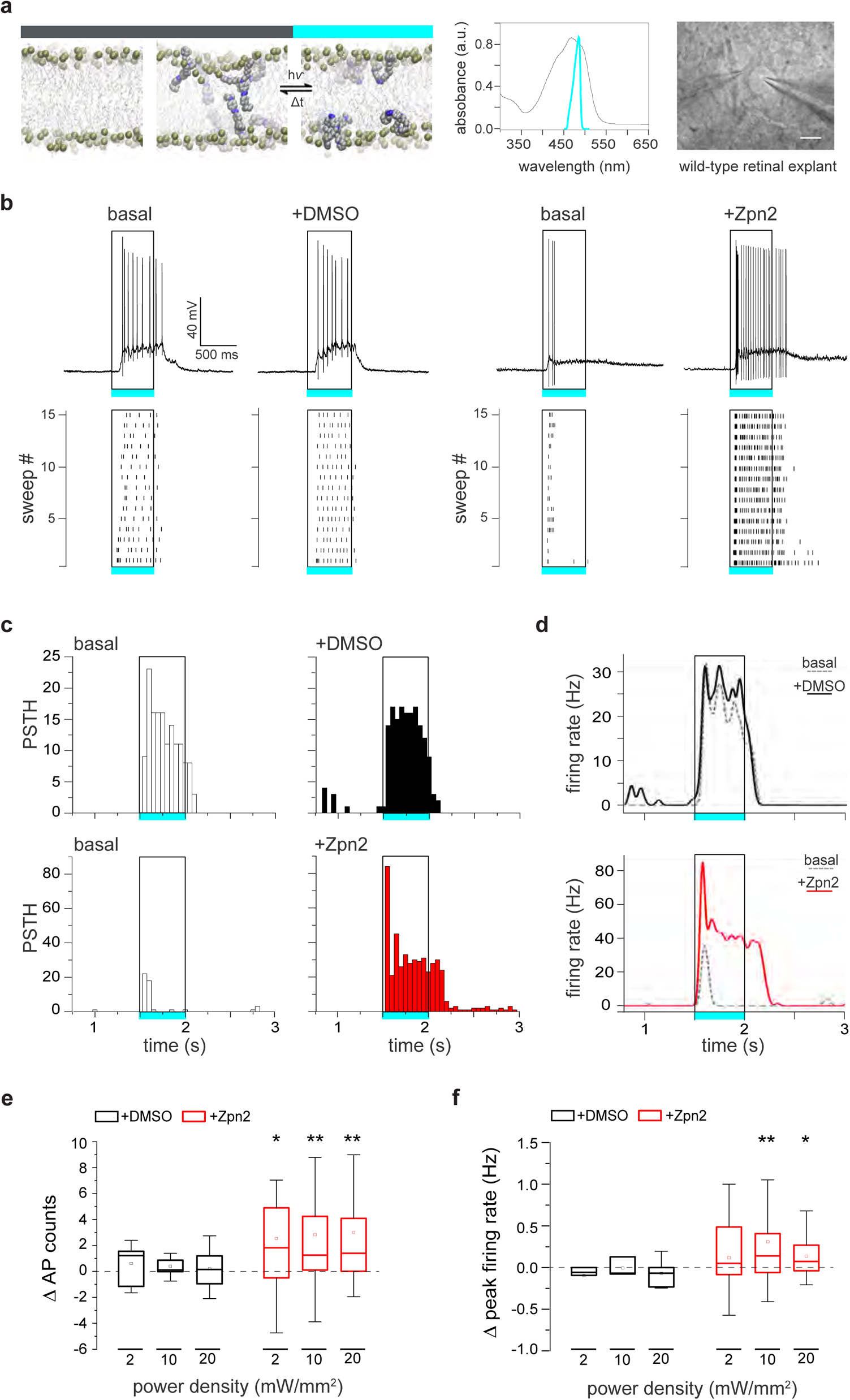
Ziapin2 enhances light evoked RGC firing activity in the WT retina. **a.** *Left:* Molecular structure of Ziapin2 and schematics of its topology after spontaneous insertion into the membrane. When Ziapin2 incorporates into a phospholipid bilayer (left snapshot) in the dark, the self-association of Ziapin2 *trans*-isomers located on opposite leaflets (middle snapshot) causes membrane shrinking, which is abolished by cyan light-induced conversion of the molecule into the *cis* conformation (right snapshot). *Middle:* Ziapin2 UV-vis absorption spectrum in DMSO (black trace; absorption peak, 470 nm), superimposed to the spectral output of the cyan LED used in the study. *Right:* Bright-field image of an RGC with a puff pipette (left) for the topic application of Ziapin2 and a patch pipette (right) for recording light-evoked firing. Scale bar, 20 µm. **b.** Representative current-clamp traces (top) and corresponding raster plots (bottom; 15 sweeps) recorded in two distinct RGCs from WT mouse retinal explants before (basal) and after the administration of either vehicle (10% v/v DMSO) or Ziapin2 (200 µM), showing an enhancement of light-evoked AP firing upon stimulation with cyan light (500 ms, 20 mW/mm^2^; open rectangle) of the Ziapin2-treated neuron. **c.** Representative peristimulus time histograms (PSTHs) of the two RGCs as in **b**, showing AP counts (50-ms bins) in response to cyan-light stimulation before (basal, open columns) and after the administration of either DMSO (top row, black columns) or Ziapin2 (bottom row, red columns). Rectangles frame the responses during the cyan light stimulus (500 ms, 20 mW/mm^2^). **d.** Changes in the firing rate of the two RGCs shown in **b** under basal conditions (dashed gray lines) and after the subsequent application of either DMSO (top, black line) or Ziapin2 (bottom, red line). The timing of the cyan-light stimulus (500 ms, 20 mW/mm^2^) is represented by the open rectangle. **e,f.** Box plots representing the cyan light-evoked changes in AP counts (**e**) and peak firing rates (**f**) with respect to the basal condition at increasing power densities (2, 10, and 20 mW/mm^2^) after the application of either DMSO (black boxes) or Ziapin2 (red boxes). Changes are expressed as the difference (Δ) between Ziapin2/DMSO puff application and the basal condition. *p<0.05, **p<0.01; two-tailed paired Wilcoxon’s signed-rank test *versus* basal (n=12 and 34 RGCs for DMSO and Ziapin, respectively). For exact p-values and source data, see the Source data file.

Here, we tested the *ex vivo* and *in vivo* ability of Ziapin2 to restore physiological light responses in degenerate retinas from the rd10 mouse bearing a recessive R560C mutation in the *Pde6b* gene essential for the proper functioning of rods ^52,53^, and the Royal College of Surgeons (RCS) rat, bearing a recessive mutation in the *Mertk* gene that impairs the ability of the retinal pigment epithelium (RPE) to phagocytose the shed outer rod and cone segments^54^. Thanks to its dual effect on intrinsic excitability, Ziapin2, by acting at the BC level, reinstated diverse light-evoked ON/OFF responses in RGCs, accompanied by the reactivation of their excitatory and inhibitory conductances. Moreover, a single intravitreal injection of Ziapin2 restored light-driven behavior and optomotor responses in blind 6-month-old rd10 mice by reactivating the distinct ON and OFF RGC populations. The results indicate that Ziapin2 is a promising molecule for reinstating visual responses at late stages of retinal degeneration through a physiological reactivation of the inner retinal processing.

## RESULTS

### Ziapin2 modulates the light-evoked activity of single RGCs in sighted animals

The bidirectional modulation of neuronal excitability by Ziapin2 prompted us to test its effects in the retina. We started with patch-clamp recordings of the light-evoked activity from single RGCs in whole-mount retinal explants isolated from sighted wild-type (WT) C56BL6/J mice (**Figure 1a**). We recorded the basal responses to full-field 500-ms flashes of either cyan (**Figure 1b**) or green (**Figure S1**) light at power densities ranging from 2 to 20 mW/mm^2^. We then approached the patched RGC with a second glass pipette for the focal application (2-min puff) of either vehicle (10% DMSO in Ames’ medium) or Ziapin2 (200 µM). Ziapin2 incorporation into the RGC membrane stimulated action potential (AP) firing in response to cyan light with respect to basal conditions, while light stimulation was ineffective in RGCs puffed with vehicle (**Figure 1b**). The modulation of cyan light-evoked firing activity by Ziapin2 was time-locked with the stimulus and consistent during the 15 sweeps of the stimulation protocol, as depicted in the respective raster plots (**Figure 1b**). Interestingly, no changes in the firing profile were detected when the same RGC was stimulated with green light falling outside the Ziapin2 absorption spectrum (**Figure S1a,b**).

Peristimulus time histogram (PSTH) analysis and firing rate computation confirmed that cyan-light stimulation reliably induced an increase in the RGC firing rate after Ziapin2 application, that was absent in vehicle-treated RGCs (**Figure 1c,d**) or in RGCs treated with Ziapin2 and subjected to green light (**Figure S1c,d**). Computation of the light-evoked changes in AP counts and firing rate showed that the effects of cyan light stimulation of Ziapin2-treated RGCs were highly significant at all the tested light power densities (2, 10, and 20 mW/mm^2^) with respect to either vehicle treatment (**Figure 1e,f**) or green light stimulation (**Figure S1e,f**).

### The changes in RGC action potential dynamics reflect the light-dependent capacitive effects of Ziapin2

To confirm the Ziapin2 photoexcitation mechanism in retinal neurons, we investigated the dynamics of AP generated in the dark or after cyan light stimulation in RGCs treated with either vehicle or Ziapin2 using the phase plane plot analysis (**Figure 2a; Figure S2**). Consistent with the capacitance increase induced by Ziapin2 in the dark, the first light-evoked AP displayed slower dynamics with respect to basal conditions, with significant decreases in the maximal depolarizing and repolarizing slopes and AP amplitude (**Figure 2a,b**). Cyan light stimulation, by phasically decreasing capacitance, brought about an opposite acceleration of AP dynamics with significant increases in the maximal depolarizing and repolarizing slopes and AP amplitude (**Figure 2a,b**). No changes in AP dynamics were observed in RGCs incubated with vehicle and subjected to the same light stimulation protocol (**Figure S2**). Similarly, no light-evoked changes in AP dynamics were observed in RGCs puffed with Ziapin2 but stimulated with green light (**Figure S3**).

**Figure 2.**
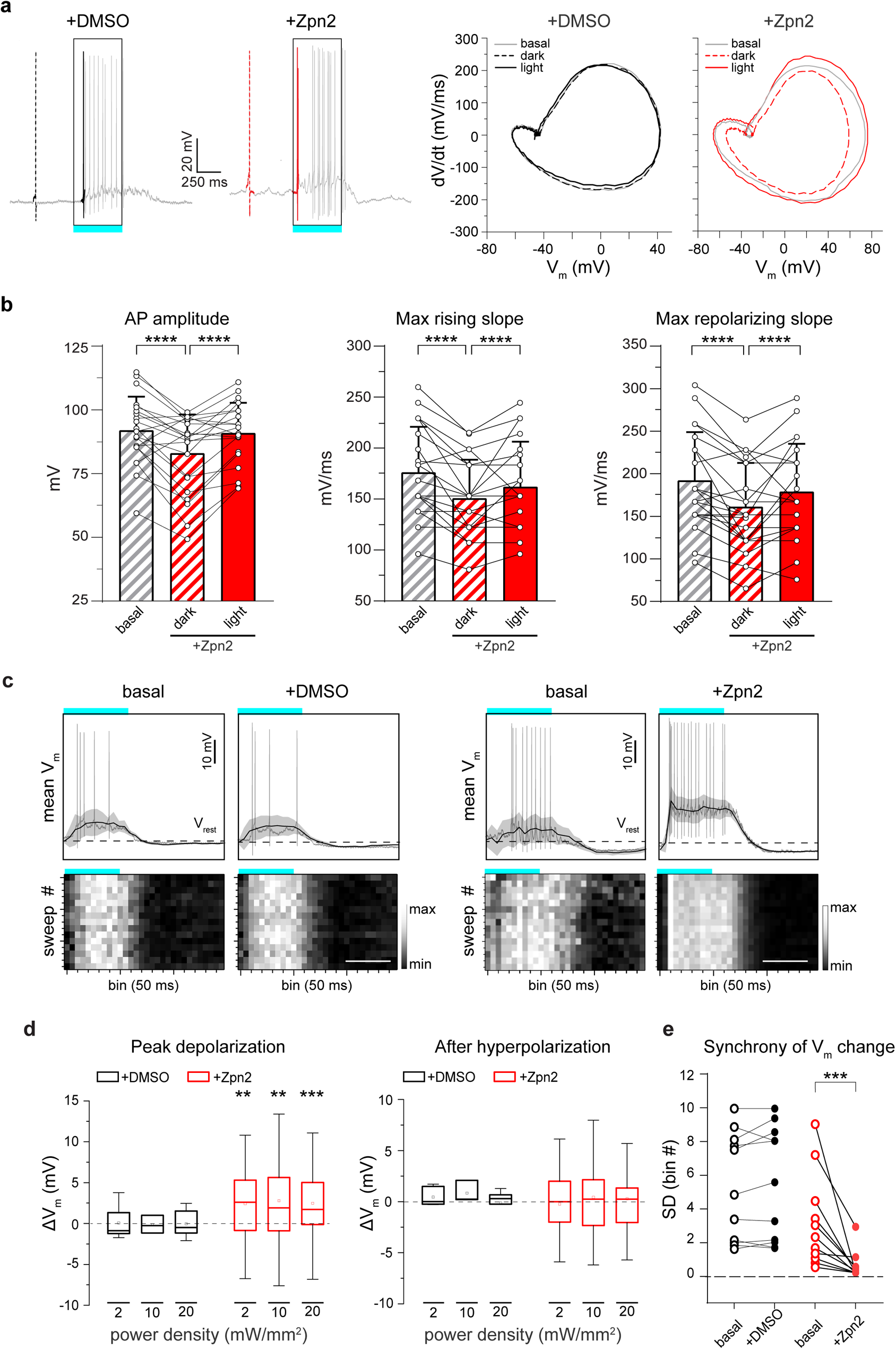
The light-evoked capacitance change by Ziapin2 induces faster AP dynamics and membrane depolarization in the WT retina. **a.** *Left:* Representative whole-cell current-clamp traces recorded in two RGCs in the presence of either vehicle (10% v/v DMSO; left, black) or Ziapin2 (200 µM; right, red) showing the first spike in the dark as a dashed line and the first light-evoked spike as a solid line. *Right:* Phase-plane plot analysis of the waveform of the first APs in the dark (dashed lines) and during cyan light stimulation (solid line; 20 mW/mm^2^, 500 ms) after puff application of either DMSO (black) or Ziapin2 (red), compared to the AP waveform in the dark under basal conditions (gray lines). The light-evoked decrease in membrane capacitance, triggered by Ziapin2 isomerization, increases AP amplitude and dynamics. **b.** Changes in AP amplitude (left), maximal rising slope (middle), and maximal repolarizing slope (right) deduced from the phase-plane plot analysis of each recorded RGC before (basal) and after Ziapin2 application in the dark and under cyan light stimulation. Bars represent means ± SEM with superimposed individual points (n = 22 RGCs). **c.** *Top:* Representative lowpass-filtered voltage traces recorded from RGCs before (basal) and after treatment with either vehicle (10% v/v DMSO; left), or Ziapin2 (200 µM; right) in a time window of 1.5 s, starting from the onset of the light stimulation (500 ms, cyan horizontal bar). The plots display the mean (± SD, gray area) V_m_ calculated within the same time window over 15 sweeps. V_rest_ (dashed line), membrane potential in the dark. APs are superimposed on the voltage traces. *Bottom:* Grayscale representation of V_m_ changes during the 15 sweeps (50-ms bins). Scale bars, 500 ms. **d.** Box plots of peak depolarization (left) and peak after-hyperpolarization (right) calculated as the difference with respect to V_rest_ (see panel **c**) in RGCs treated with either vehicle (10% v/v DMSO; black) or Ziapin2 (200 µM; red) and stimulated with cyan light at increasing power densities (2, 10, and 20 mW/mm^2^). Ziapin2 significantly enhances membrane depolarization in response to light stimulation, while it does not affect the extent of after-hyperpolarization (n=12 and 34 for DMSO and Ziapin2 puffs, respectively). **e.** Evaluation of the synchrony of voltage changes in response to cyan light before (basal) and after Ziapin2 or DMSO puff application. Synchrony was calculated, for each recorded RGC, as the standard deviation of the bin experiencing the maximum voltage change in each sweep (n=10 and 11 RGCs for DMSO and Ziapin2 puffs, respectively). **p<0.01, ***p<0.001, ****p<0.0001; two-tailed paired Wilcoxon’s signed-rank *versus* basal. For exact p-values and source data, see the Source data file.

We then filtered the recordings with a lowpass filter to extract the slow subthreshold membrane potential oscillations in a time window of 1.5 s after the onset of the light stimulus (**Figure 2c**). We observed a significant increase of the peak membrane depolarization during the light response at all tested power densities, while the peak afterhyperpolarization was not affected (**Figure 2d**). Ziapin2 focused the onset of depolarization and the return to basal V_m_ levels exactly at the switching on/off of the light stimulus, while RGCs under basal conditions or treated with vehicle displayed a much wider temporal variability in V_m_ changes. To quantitatively evaluate this effect, we calculated, for each recorded RGC, the standard deviation of the bins in which the maximum V_m_ changes occurred over the 15 sweeps. The results show that indeed the presence of Ziapin2 in the RGC membrane significantly synchronizes sub-threshold light-evoked membrane potential oscillations, reducing response variability (**Figure 2e**).

### Ziapin2 restores the natural encoding capabilities of RGCs in retinal explants from blind rd10 mice

Given the dark/light bifunctional activity of Ziapin2 demonstrated in spiking RGCs of sighted retinas, we sought to test whether it could restore the light sensitivity and the complex processing of the retinal network in light-insensitive degenerate retinas. To test this hypothesis, we conducted patch-clamp experiments on blind retinas taken from 6-month-old rd10 mice. In a first attempt, we puffed Ziapin2 (200 µM) directly on the recorded RGCs, as done with normal sighted retinal explants (**Figure S4a**). However, in RGCs from dystrophic retinas, cyan light stimuli did not evoke APs but only light-dependent subthreshold depolarizations that were absent in response to green light (**Figure S4b,c**). This indicates that, in light-insensitive retinas, Ziapin2 was unable to make RGCs reach the firing threshold in the absence of concomitant light-evoked synaptic inputs from second-order retinal neurons. Thus, to permeate with Ziapin2 all retinal neurons spared by degeneration, we immersed retinal explants in a solution containing Ziapin2 (10 µM for 30 min) before transferring them to the recording chamber and evaluating the Ziapin2-specific effects to cyan *versus* green light stimuli. Under these conditions, RGCs fired trains of APs in response to cyan, but not green light (**Figure S4d**).

Indeed, after whole retina treatment, Ziapin2 reactivated a variety of RGC responses, including ON and OFF responses to cyan light with varying dynamics, ranging from transient to sustained and suppressed (**Figure 3a-d**). Total AP counts and peak instantaneous firing rate significantly increased in response to cyan light at all tested power densities, ranging from 2 to 30 mW/mm^2^, but not to green light (**Figure 3e**). When we retrospectively studied the 3D morphology of the recorded RGCs labeled with AlexaFluor-633 by confocal microscopy, as previously described ^55^, we found that the responses evoked by cyan light were always consistent with the stratification of the dendrites in the inner plexiform layer of all recorded RGCs (**Figure 3a**).

**Figure 3.**
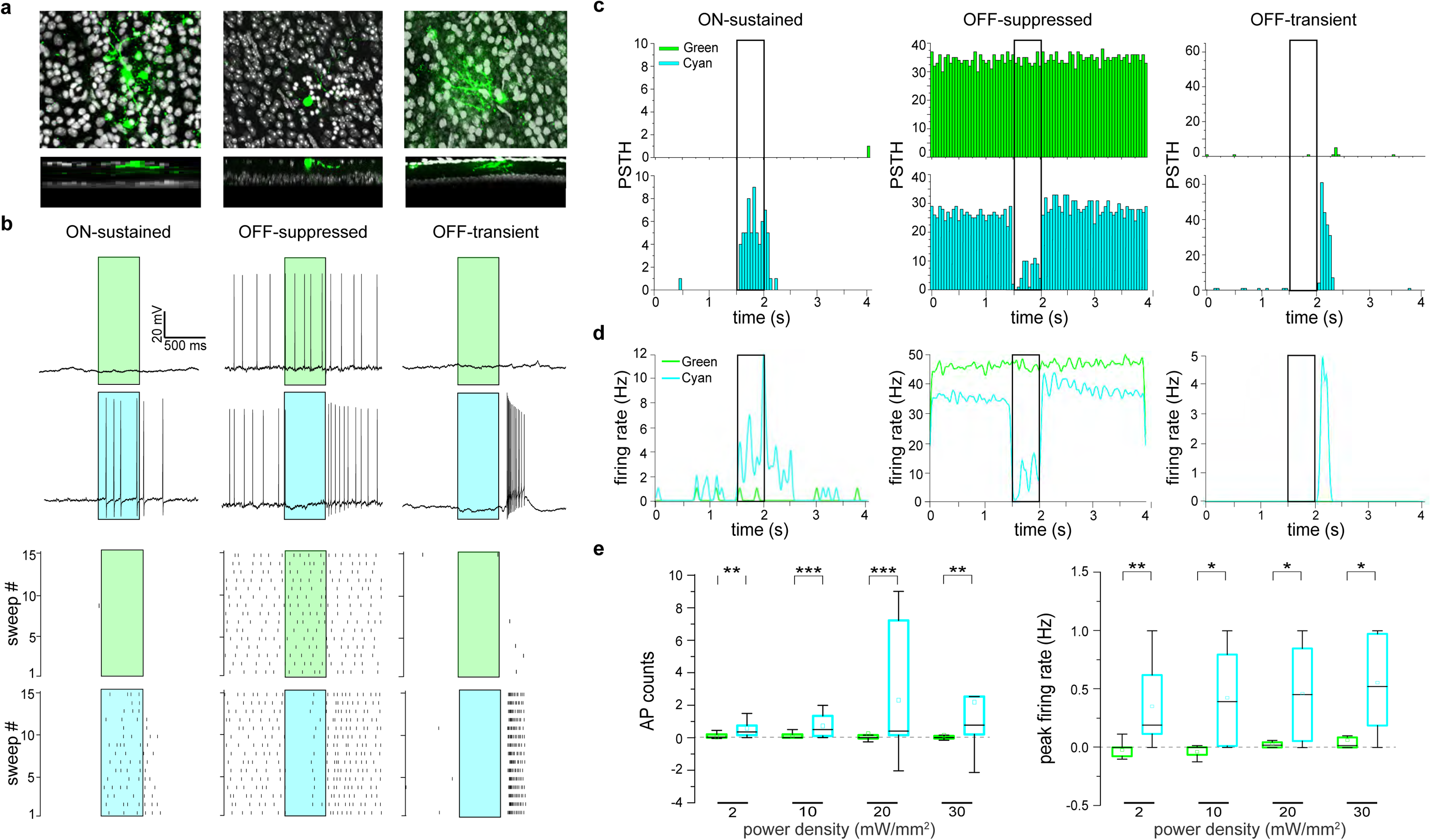
Ziapin2 restores physiological responses to light in RGCs from degenerate rd10 retinas. **a.** *Top:* Morphological reconstructions of the RGCs recorded in **b** filled with Alexa Fluor 633 (green). *Bottom:* Transversal sections of the same Z-projections, showing the dendritic tree of the recorded RGCs in distict IPL *sublaminae*. The morphology and dendritic stratification of the three RGCs correspond to their ON-sustained (left), OFF-suppressed (middle), and OFF-transient (right) physiological responses to light. **b.** *Top:* Representative responses of the RGCs shown in **a** to full-field illumination with either green or cyan light (500 ms; 20 mW/mm^2^) after incubation of the blind retinal explants in the presence of Ziapin2 (10 µM). No light-dependent firing activity was detected upon illumination with green light, while cyan light stimulation induced various responses resembling those of sighted animals. *Bottom:* Raster plots show firing activity of the same RGCs recorded over 15 consecutive sweeps. Light stimuli are shown as cyan- or green-shaded areas. **c,d.** Representative PSTHs (**c**) and instantaneous firing rate (**d**) of the RGCs recorded in **b**. Bars represent AP counts (50-ms bins) recorded in response to either green (top panels) or cyan (bottom panels) stimulation. Light stimuli (500 ms) are shown as open rectangles. No modulation of firing activity is evoked in response to green light. **e.** Box plots of the light-evoked AP counts (*left*) and peak firing rate (*right*) in response to 500-ms stimulation with either green or cyan light (colored boxes) in a time window starting from the onset to 1 s after the end of the light stimulus and normalized to the spontaneous activity recorded in the dark. Light stimulation was applied at increasing power densities (2, 10, 20, and 30 mW/mm^2^). *p<0.05, **p<0.01, ***p<0.001; two-tailed paired-sample Wilcoxon’s signed-rank test (n=14). For exact p-values and source data, see the Source data file.

To explore the complexity of RGC responses that might arise in Ziapin2-treated blind retinas, we recorded light-evoked responses using high-density multielectrode arrays (HD-MEA) that allow to simultaneously follow hundreds of RGCs ^56,57^. Green light stimuli did not substantially affect the overall RGC firing (**Figure 4a,b**), except for the typical slow, sustained responses of a few intrinsically photosensitive RGCs (shown in the left raster plot of **Figure 4b**). On the contrary, cyan light stimuli recreated the plethora of responses typically recorded in normally sighted retinas: brisk to sluggish ON-sustained and ON-transient cells, ON-OFF cells, and OFF-suppressed or OFF-transient cells could be identified (**Figure 4c**). Notably, Cell 4 shows a clear sign of adaptation, meaning that Ziapin2 restored subtle but fundamental network mechanisms.

**Figure 4.**
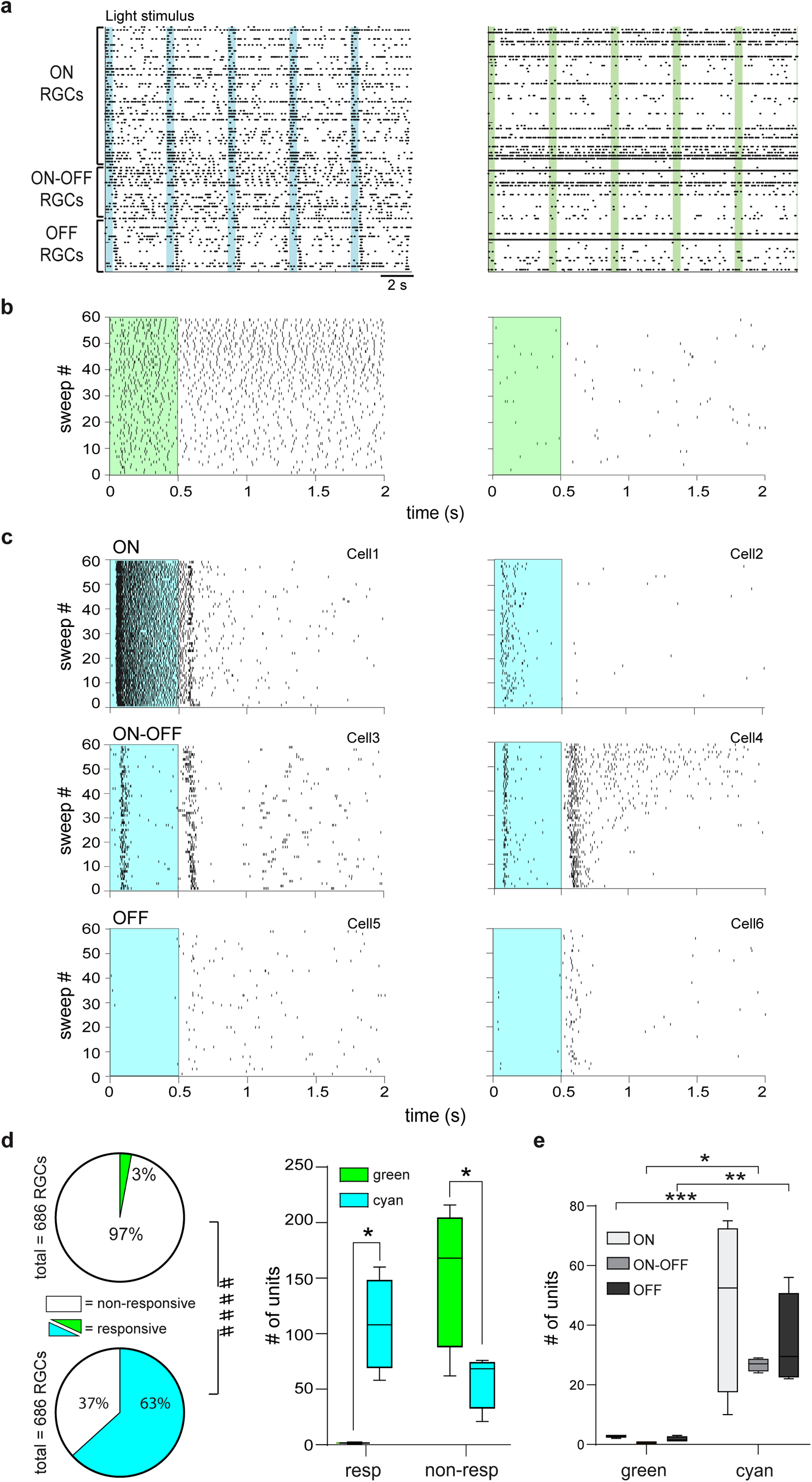
Ziapin2 restores the ability of degenerate rd10 retinas to analyze visual information in segregated channels. **a.** Representative AP raster plots obtained from HD-MEA recordings of the same rd10 blind retinal explant after incubation with Ziapin2 (10 µM) stimulated with either green (left) or cyan (right) light (500 ms; 2 mW/mm^2^). Each row represents the firing activity of single RGCs that were sorted for polarity. Light-dependent firing activity was only restored with full-field cyan, but not green, light stimulation. Light pulses are indicated with vertical color-coded bars. **b.** Representative raster plots of APs generated by two RGCs with very different spontaneous activity over 60 sweeps. In both cells, green light stimulation (green rectangle) could not modulate firing activity. **c.** Representative raster plots of AP firing of different classes of RGCs in response to full-field stimulation with cyan light (cyan rectangle). Cyan stimuli evoked ON-sustained (Cell1), ON transient (Cell2), ON-OFF (Cell3, Cell4), OFF-suppressed (Cell5) and OFF-transient (Cell6) responses. **d.** *Left:* Pie charts represent the percentage of RGCs responsive to light stimulation with green (top) and cyan (bottom) light with respect to the total number of RGCs recorded by HD-MEA (numbers reported in parenthesis). ***p<0.001, Fisher’s exact test; responsive/non-responsive RGCs to green *versus* cyan light. *Right:* Box plots showing the number of responsive and non-responsive RGCs to either cyan or green light stimuli in the retinas from n=4 rd10 mice. **e.** Box plots showing the number of ON, ON-OFF, and OFF RGCs responsive to either cyan or green light stimuli in the retinas from n=4 rd10 mice. ^####^p<0.0001, Fisher’s exact test; responsive/non-responsive RGCs to green *versus* cyan light; *p<0.05, **p<0.01, ***p<0.001, two-way ANOVA/Holm-Šídák tests (n=4). For exact p-values and source data, see the Source data file.

Using HD-MEA, we also evaluated the percentage of light-responsive RGCs reactivated by Ziapin2 in the degenerate retinas by considering responsive those RGCs exhibiting light-evoked changes in firing rate larger than the mean spontaneous firing rate plus 2 x SD. While only 3% of the RGCs in all the recorded retinas were responsive to green light stimulation, a significantly higher percentage of cells (63%) were responsive to cyan light in Ziapin2-treated retinas (Fisher’s exact test, p<0.0001; **Figure 4d**). When the results of the individual HD-MEA experiments were analyzed, we observed that Ziapin2 was able to significantly restore the responses to cyan light in a variety of RGC subclasses, as compared to green light that was practically ineffective (**Figure 4d,e**).

Being the capacitance change transient in nature, the Ziapin2 effects have the potential to respond to high frequency stimuli. Thus, we asked how frequently the light-induced Ziapin2 stimulation could be repeated to boost cell excitability. By subjecting blind rd10 explants to progressively shorter cyan light flashes at increasing frequency, as compared to the single 250-ms light flash at 0.25 Hz, we found that the Ziapin2-induced increase in the RCG firing rate was unaffected up to 10 Hz light stimulation, decreased but still present at 15 Hz, and practically absent at 30 Hz (**Figure S5**).

Although the main objective was to characterize the effect of Ziapin2 on the denervated BCs, we asked whether the Ziapin2 effect was specific to degenerating retinas or whether it was also present in the whole circuitry of WT retinas. We incubated WT retinal explants in either vehicle (0.5% DMSO) or Ziapin2 (10 µM) solution and recorded the responses to cyan light of RGC populations by HD-MEAs. We found that Ziapin2-treated retinas displayed a similar number of responsive ON, OFF, ON/OFF RGCs with sustained or transient firing pattern (**Figure S6a,b**), but that the firing rate of sustained ON and OFF RGCs in response to cyan light was greatly and significantly enhanced with respect to vehicle-treated WT retinas (**Figure S6c**).

### Ziapin2 accelerates action potential dynamics in RGCs from degenerate rd10 retinas

Blind retinas undergo significant and profound changes in the anatomical and functional network organization concomitant with photoreceptor loss, resulting in the impaired ability of RGCs to receive light-evoked synaptic inputs and trigger AP firing to process visual information.

We firstly verified that the evoked firing in degenerate retinas was indeed triggered by the sudden decrease in membrane capacitance by Ziapin2 *trans*-to-*cis* isomerization. To this aim, individual RGCs were locally puffed with Ziapin2 (200 µM) and stimulated with either cyan or green light (**Figure S7a**). In agreement with our previous observations in primary hippocampal neurons ^49^, cyan, but not green, light stimuli significantly decreased RGC capacitance, without affecting membrane resistance, resulting in a light-dependent decrease of the membrane time constant (**Figure S7b,c**). To gain further insights into Ziapin2 capacitive effects on the passive properties of RGCs in rd10 retinal explants, we analyzed changes in AP dynamics by phase plane plot analysis. Retinas were incubated with Ziapin2 (10 µM for 30 min) and interrogated with cyan light to activate Ziapin2 or green light as a negative control. In the dark or in response to green light, no differences were observed in the firing pattern and AP waveforms (maximum AP peak values and maximum rising and repolarizing slopes; **Figure 5a,b**). On the contrary, stimulation with cyan light caused a noticeable acceleration of AP kinetics with significant increases in peak amplitude and faster rising and repolarizing phases with respect to AP generated by the same cells in response to green light stimulation (**Figure 5c**). When we applied a lowpass filter to cut out APs and detect slow voltage oscillations (**Figure 5d**), we could uncover a significant increase in the amplitudes of both peak depolarization and afterhyperpolarization in response to cyan, but not green, stimuli at all tested power densities (2-30 mW/mm^2^; **Figure 5e**).

**Figure 5.**
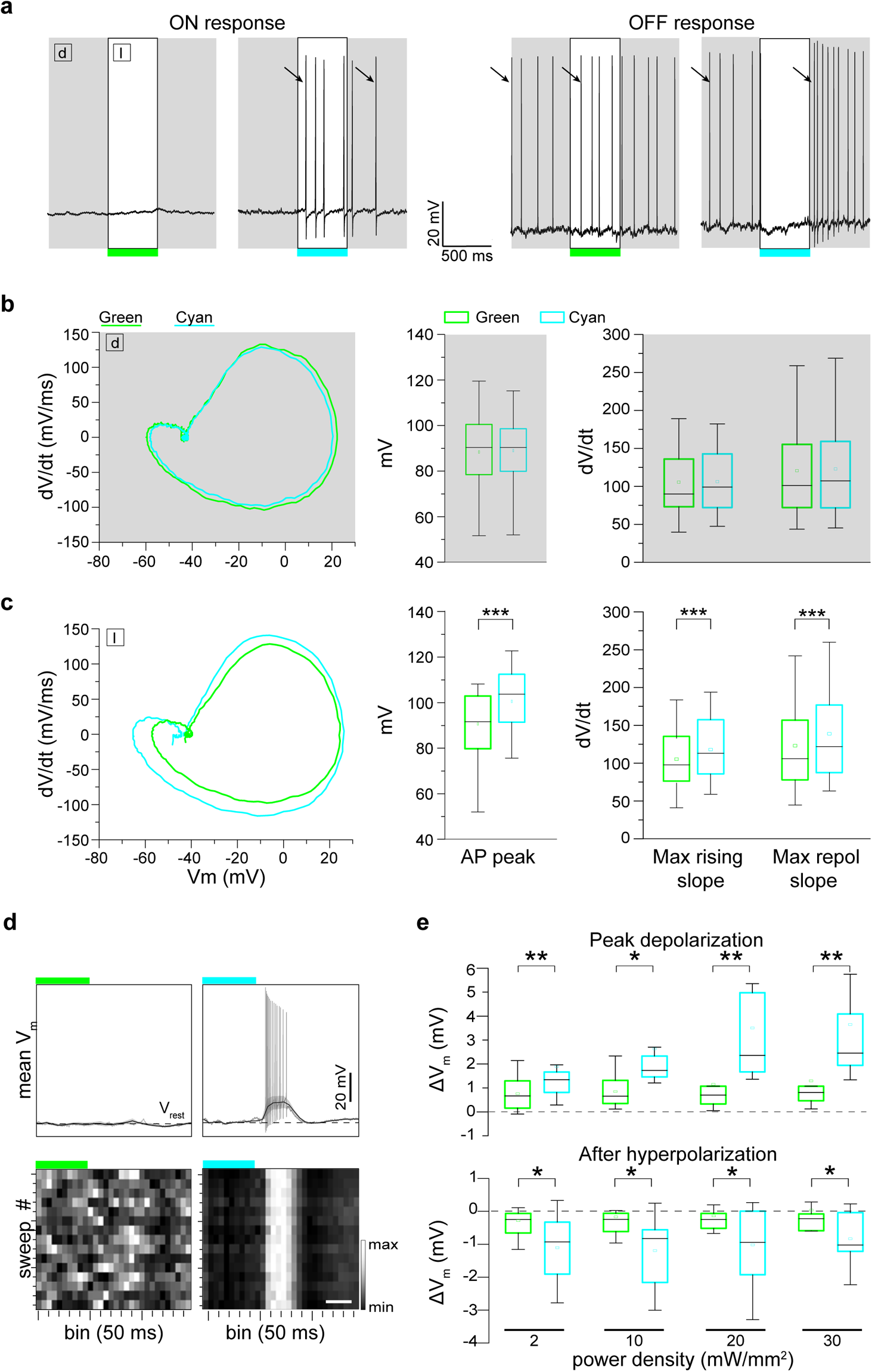
The light-evoked capacitance change by Ziapin2 induces faster AP dynamics and membrane in degenerate rd10 retinas. **a.** Representative current-clamp traces, recorded in two RGCs of blind rd10 retinas incubated with Ziapin2 (10 µM), showing ON (left) and OFF (right) responses to cyan light (right traces), but not to green light (left traces). The open rectangles represent the timing of the light stimuli (500 ms, 20 mW/mm^2^). Arrows indicate the first AP in the dark (d) and in response to light (l) used to perform the waveform analysis shown in **b**. **b.** *Left:* Representative phase-plane plot analysis of the first AP generated in the same RGCs shown in **a** in the dark (d), before light stimulation. *Right:* Box plots showing AP peak, maximal rising slope, and maximal repolarizing slope in the dark calculated from the phase plot analysis of all recorded RGCs **c.** *Left:* Representative phase-plane plot analysis of the first AP generated in the same RGCs shown in **a** during light stimulation (l) with either green or cyan light. *Right:* Box plots of AP peak, maximal rising slope, and maximal repolarizing slope after stimulation with either green or cyan light. The cyan light-induced decrease in membrane capacitance by Ziapin2 increases the AP amplitude and dynamics with respect to green light stimulation. ***p<0.001; 2-tailed paired Wilcoxon’s signed-rank test (n=28). **d.** *Top:* Representative lowpass-filtered voltage traces recorded from RGCs incubated with Ziapin2 (10 µM) in a time window of 1.5 s, starting from the onset of either green (left) or cyan (right) light stimulation (500 ms, 20 mW/mm^2^). The plots display the mean (± SD, gray area) V_m_ calculated within the same time window over 15 sweeps. V_rest_ (dashed line), membrane potential in the dark. APs evoked with cyan light are superimposed on the voltage traces. *Bottom:* Grayscale representation of V_m_ changes during the 15 sweeps (50-ms bins). Scale bar, 250 ms. **e.** Box plots of peak depolarization (*top*) and peak after-hyperpolarization (*bottom*) calculated as the difference with respect to V_rest_ (see panel **d**) in RGCs treated with Ziapin2 and stimulated with either green or cyan light (colored boxes) at increasing power densities (2, 10, 20 and 30 mW/mm^2^). *p<0.05; **p<0.01; two-tailed paired Wilcoxon’s signed-rank test cyan-*versus* green-light stimulation (n=14). For exact p-values and source data, see the Source data file.

### Ziapin2 restores light-evoked excitatory and inhibitory synaptic inputs to RGCs in degenerate rd10 retinas

We showed that Ziapin2 restores the ability of degenerate retinas to reactivate segregated channels of visual information processing, but only when the photoswitch was applied to all retinal neurons and not only to a single RGC (see Figure S4). These findings raise the question of whether the variety of light evoked RGC responses results from the restoration of the entire retinal network functionality, rather than from a direct effect on the intrinsic excitability of RGCs. To answer this question, we performed patch-clamp experiments in voltage-clamp configuration to measure light-evoked synaptic inputs to RGCs in blind rd10 retinas incubated with Ziapin2 (10 µM for 30 min). Before accessing the intracellular compartment, we assessed the polarity of the RGC by recording cyan light-evoked spiking activity in cell-attached configuration, using green light as negative control (see the representative OFF RGC in **Figure 6a**). Interestingly, once the whole-cell configuration was reached, we found out that APs were generated by the activation of light-evoked synaptic conductances in RGCs, which were negligible with green light stimulation (**Figure 6b**). Indeed, Ziapin2 restored both excitatory and inhibitory conductances (**Figure 6c**), with cell polarity and AP generation resulting from their fine temporal interplay. Overall, Ziapin2 activation in retinal neurons of blind retinal explants significantly reactivated physiological excitatory (g_e_) and inhibitory (g_i_) synaptic inputs to RGCs at both 2 and 20 mW/mm^2^ power densities (**Figure 6d**).

**Figure 6.**
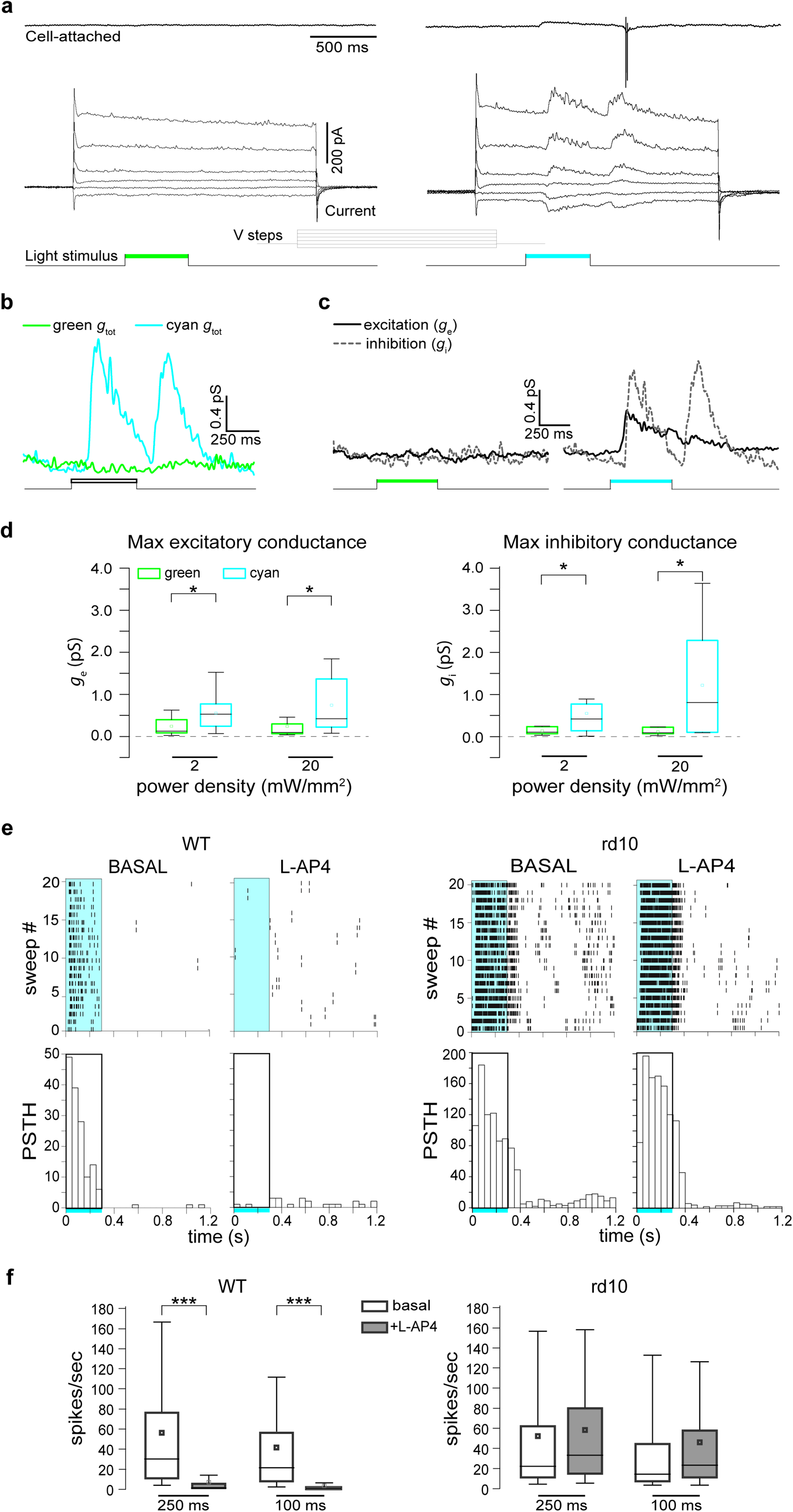
Ziapin2 restores the light-evoked excitatory and inhibitory synaptic inputs to RGCs in degenerate rd10 retinas. **a.** *Top:* Representative voltage-clamp recordings in the cell-attached configuration of an OFF-RGC from a blind rd10 retina after 30 min incubation with Ziapin2 (10 µM) and subjected to full-field stimulation with either green (left) or cyan (right) light (500 ms, 20 mW/mm^2^). *Bottom:* Light-induced currents were recorded in the same RGC after reaching the whole-cell configuration and applying voltage steps from -86 to 34 mV. Light-evoked inward and outward currents were recorded only in response to cyan light stimulation (right panel), eventually triggering the AP recorded in cell-attached configuration. The light stimuli are shown as color-coded bars in the bottom traces. **b.** Total conductances in the same Ziapin2-incubated RGC shown in **a** in response to either green or cyan light stimulation calculated from the current traces. The stimulation with cyan, but not with green light, induces a marked activation of RGC synaptic conductances. **c.** The excitatory (*g*_e_) and inhibitory (*g*_i_) conductances evoked, in the same RGC shown in **a**, by either green (left) or cyan (right) light stimuli were extrapolated from the voltage-clamp recordings. Ziapin2 restores excitatory and inhibitory RGC synaptic conductances in response to cyan light. **d.** Box plots of the maximum values of excitatory (*g*_e_; *left*) and inhibitory (*g*_i_; *right*) conductances calculated from all RGCs stimulated with green (green boxes) and cyan (cyan boxes) light at 2 and 20 mW/mm^2^ power densities. *p<0.05; 2-tailed paired Wilcoxon’s signed-rank test, cyan-*versus* green-light stimulation (n=12). **e.** *Left:* Representative HD-MEA recordings (raster plots of 20 sweeps and respective PSTHs shown below) of an ON-RGC from a healthy WT retina stimulated with cyan light (2 mW/mm^2^; 250 ms; rectangle) under basal conditions and after the application of the mGluR6 agonist L-AP4, which abolishes the light-evoked activation of rod and cone-ON BCs. *Right:* The same protocol was applied to a representative RGC from blind rd10 retinas incubated with Ziapin2 (10 µM) and stimulated with cyan light before (basal) and after the application of L-AP4. The lack of L-AP4 effect indicates that the reactivation of RCGs in degenerate retinas by Ziapin2 occurs downstream of the photoreceptor/BC synapses. **f.** Box plots of the RGC firing activity in healthy WT retinas (*left*) and Ziapin2-treated blind rd10 retinas (*right*) stimulated with cyan light (2 mW/mm^2^) for 100 and 250 ms in the absence or presence of L-AP4 (n=322 and 266 RGCs for WT and rd10 retinas, respectively from n=2 WT and n=2 rd10 mice). ***p<0.001, two-tailed paired Wilcoxon’s signed-rank test, basal *versus* L-AP4. For exact p-values and source data, see the Source data file.

To further investigate the contribution of synaptic transmission in the generation of Ziapin2-mediated responses, we perfused WT and rd10 retinal explants with the mGluR6 agonist L-AP4 to prevent photoreceptor-mediated signal transmission to second-order neurons ^58^ and analyzed the RGC firing responses to cyan light by HD-MEA recordings. Strikingly, in representative RGCs, the firing responses to 250-ms cyan light stimuli were totally abolished by L-AP4 in WT retinas, while they were virtually unaffected by the drug in Ziapin2-treated rd10 retinas (**Figure 6e**). Statistical analysis of the firing responses of RGC populations to 100- and 250-ms cyan stimuli obtained in independent experiments confirmed the highly significant inhibition of ON firing responses by L-AP4 in WT retinas, but not in Ziapin2-treated rd10 retinas (**Figure 6f**). This result rules out the possibility that the activation of residual photoreceptors mediates the light-evoked currents and APs recorded in RGCs of rd10 retinas, indicating that Ziapin2 acts primarily at the level of BCs.

To verify whether the reactivation of OFF responses following cyan stimuli in Ziapin2-treated rd10 retinas also involved rod BCs and AII amacrine cells, we perfused retinal explants with the glycine receptor blocker strychnine (10 µM; **Figure S8a**). We found that strychnine significantly decreased the percentage of OFF and ON/OFF RGCs by ∼25%, while no significant effects were observed on the percentage of ON-RGCs (**Figure S8b**) and in the relative prevalence of responding RGC subtypes (**Figure S8c**).

### The reactivation of retinal processing in degenerate retinas by Ziapin2 is independent of species and gene mutation

*Retinitis pigmentosa* is a monogenic disease resulting from over a hundred distinct mutations hitting genes expressed in either photoreceptors or RPE cells. To assess whether the reactivation potential of Ziapin2 in degenerate retinas is independent of the specific mutation and dysfunctional cell type, we explored Ziapin2 effects on the retinas from 10-month-old blind RCS rats, an animal model of rod-cone degeneration due to mutation of the *Mertk* gene that causes dysfunction of the RPE. Fully degenerate retinas explanted from RCS rats were incubated with Ziapin2 (10 µM for 30 min) and assessed for light sensitivity by patch-clamp recordings of RGCs in response to either cyan light or green light as a negative control.

Notwithstanding the different animal species and gene mutation, we were able to elicit light-evoked responses similarly to those observed in Ziapin2-treated blind rd10 retinas, with the presence of ON-sustained, OFF-suppressed and OFF-transient responses, which were not detectable when the same RGCs were stimulated with green light (**Figure S9a**). PSTH analysis clearly showed that the increase in AP firing was detected within 1.5 sec time-window from the onset of the light stimulus exclusively upon illumination with cyan light, but not with green light, testifying that Ziapin2 was responsible for the effects (**Figure S9b**). The peak rate of AP firing was measured either during the cyan light stimulation or immediately after the end of the stimulus, mimicking the well-known ON- and OFF-responses of WT RGCs (**Figure S9c**). Finally, we found significant increases in both light-evoked AP counts and peak firing rate in all recorded RGCs in response to cyan light stimulation at all tested power densities (2-30 mW/mm^2^), in the absence of responses to green light (**Figure S9d,e**).

We next characterized these responses of RGCs with the AMPA/kainate receptor antagonist CNQX (10 µM) ^59^ and the mGluR6 agonist L-AP4 (20 µM) ^58^. CNQX entirely abolished OFF-responses evoked by cyan light in Ziapin2-treated RCS retinas, while L-AP4 did not affect the ON response to cyan light (**Figure S10a**). We also analyzed the RGC subthreshold membrane voltage oscillations, as shown in Figure 2, and observed that the Ziapin2-dependent depolarization in response to cyan light was virtually unaffected by L-AP4 and was decreased, but not abolished, by CNQX (**Figures S10b**). These results further confirm that the Ziapin2-evoked depolarization of synaptically isolated RGCs in blind retinas is insufficient to reach the firing threshold and that the light-evoked RGC firing in blind retinas results from restoring the fine interplay between second-order neurons and interneurons of the inner retina.

We next focused on the effects of CNQX and L-AP4 application on light-evoked synaptic inputs to RGCs in the RCS blind retina. CNQX notably abolished the light-evoked synaptic inputs to RGC, supporting the hypothesis that Ziapin2 reinstates light-evoked responses in blind retinas by acting on the entire retinal network (**Figure S10c**, *left panel*). This was further confirmed by the fact that the synaptic currents generated by cyan light stimulation in RGCs were superimposable before and after application of L-AP4, indicating that signal transmission does not rely on photoreceptors activation (**Figure S10c**, *right panel*).

Notably, recordings performed during stimulation of the same RGC with green light revealed the absence of light-evoked responses, as well as negligible effects of the two drugs on RGC firing activity (**Figure S11a**). Moreover, CNQX and L-AP4 did not elicit any detectable effects on membrane voltage and basal RGC conductances measured during light stimulation (**Figure S11b,c**).

To corroborate this hypothesis, healthy retinas from age-matched congenic RDY rats and Ziapin2-treated degenerate retinas from dystrophic RCS rats were recorded in parallel on HD-MEA before and after the application of L-AP4 (**Figure S12**). In normally sighted RDY retinas, light evoked ON responses from ON- and ON-OFF RGCs were completely abolished, while the OFF response of the ON-OFF RGC was unaffected (**Figure S12a,c**). On the opposite, dystrophic RCS retinas incubated with Ziapin2 did not show any alteration of the cyan light-evoked RGC firing profile after application of L-AP4 for both ON and ON-OFF RGCs (**Figure S12b,d**). Statistical analysis of the firing responses of RGCs to cyan light stimuli obtained in independent experiments confirmed that L-AP4 significantly abolished the RGC firing response only in sighted RDY RGCs, while it was totally ineffective in blind RCS RGCs (**Figure S12e**).

### *In vivo* intravitreal injection of Ziapin2 restores visually driven behavior in aged blind rd10 mice

To address the possibility that Ziapin2 can act as a potential therapeutic tool for vision restoration *in vivo*, we intravitreally injected it (200 µM/10% final vitreal concentration of DMSO) in 6-month-old blind rd10 mice. Age-matched WT and blind rd10 mice were both injected with the vehicle and used as controls. During the two weeks after the injection, mice were studied for their performance in light-driven behavioral tests (**Figure 7a**). The visual rescue of blind mice was evaluated in the Dark-Light Box, which is based on the innate aversion of rodents to illuminated open spaces generating anxiety ^60^ (see Materials and Methods). Unconstrained mice could freely move in the whole apparatus, and the time spent in each chamber was measured for 3 min under dark conditions, followed by 3 min with a white 5 lux-dim light shining in the left chamber. In the total dark test, the WT mice and vehicle-treated and Ziapin2-treated blind rd10 mice behaved very similarly, as evaluated from both the time spent in the left chamber and in its open center area (**Figure 7b,c**). Conversely, when light was switched on, mice intravitreally injected with Ziapin2 recovered their light-avoidance behavior, significantly decreasing both the time spent in the illuminated chamber and in its open center area with respect to vehicle-injected blind controls and approaching the performances of WT mice (**Figure 7d,e**). Notably, analysis of the time-course of the effects up to 2 weeks after the injection showed that the improvement in light-driven behavior induced by Ziapin2 was persistent over time, as also demonstrated by the individual performances of the animals (**Figure S13a,b**).

**Figure 7.**
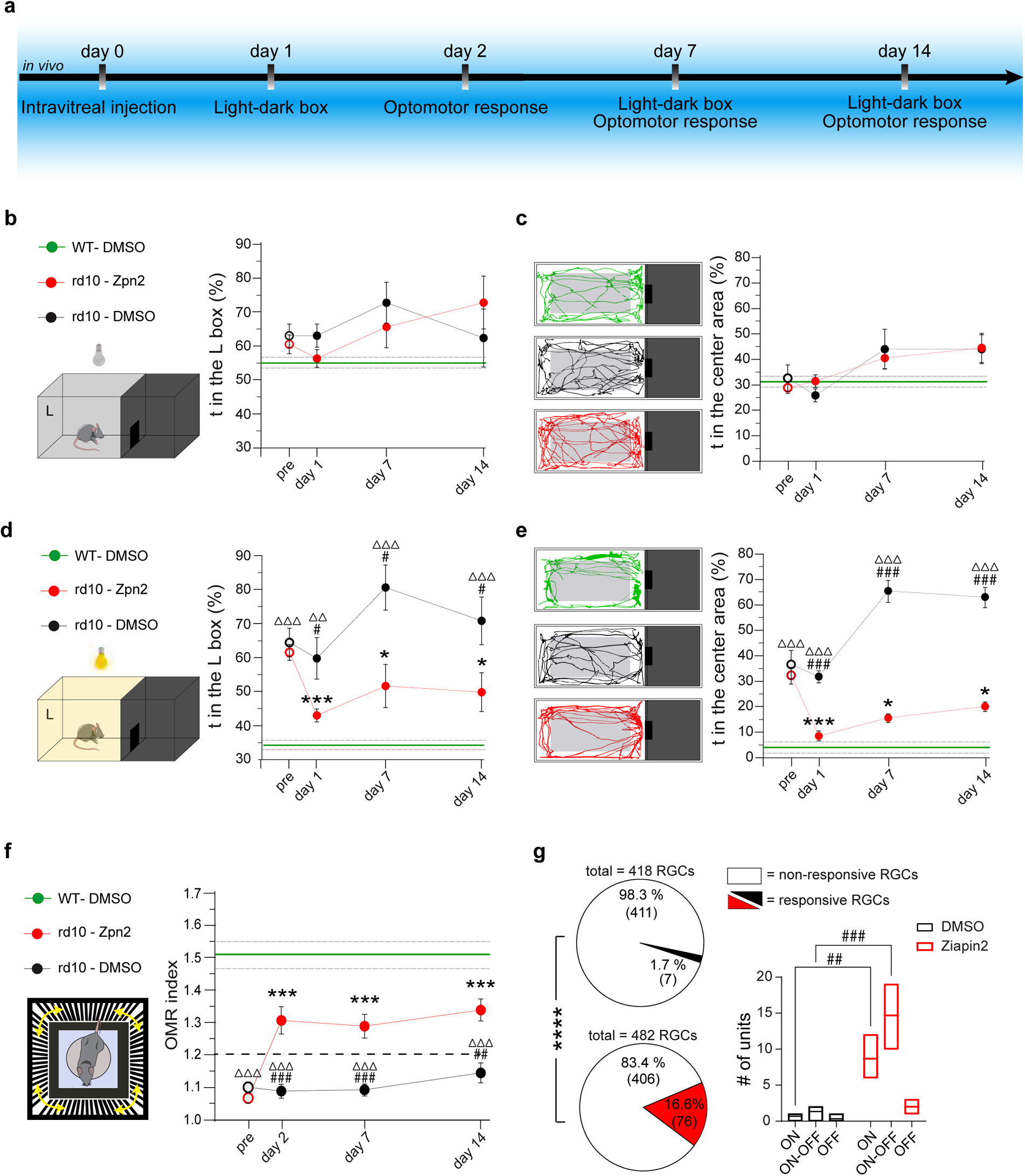
Visually driven behaviors and physiological RGC discharges are restored in blind rd10 mice after *in vivo* administration of Ziapin2. **a.** Experimental timeline indicating the *in vivo* procedures performed in 6-month-old blind rd10 mice and aged-matched WT mice subjected to the bilateral intravitreal injection of either vehicle (10% DMSO final vitreal concentration) or Ziapin2 (200 µM final vitreal concentration) in saline. **b.** Schematic representation of the Light-Dark box apparatus kept entirely in the dark. The time-course on the right shows the mean (± SEM) percent time spent in the left compartment (L) over the total duration of the test by blind rd10 mice injected with either vehicle (DMSO; black) or Ziapin2 (red). The mean (± SEM) value of age-matched, vehicle-injected WT mice (green) over the two-weeks of observation is shown for comparison (mean: solid horizontal line; SEM, broken lines). Animals did not show any significant preference between the two compartments. **c.** Representative animal tracks in the left compartment for the three experimental groups described in **b**. The time-course on the right shows the mean (± SEM) percent time spent in the center area of the left compartment (gray rectangle) over the 3-min of observation. The mean (± SEM) value of vehicle-injected WT mice (green) over the two-weeks of observation is shown for comparison. **d.** Schematic representation of the Light-Dark box apparatus during the light experimental condition (5-lux white light in the left compartment). The time-course on the right shows the mean (± SEM) percent time spent in the lighted compartment over the 3-min test by the three experimental groups. The mean (± SEM) value of age-matched, vehicle-injected WT mice (green) over the two-weeks of observation is shown for comparison. **e.** Representative animal tracks in the lighted left compartment for the three experimental groups described in **d**. The time-course on the right shows the mean (± SEM) percent time spent in the center area of the left compartment. The mean (± SEM) value of vehicle-injected WT mice (green) over the two-weeks of observation is shown for comparison. **f.** *Left:* Schematic representation of the Optomotor Response (OMR) apparatus. The unconstrained mouse instinctively follows the grating patterns rotating around it (yellow arrows) with synchronized head movements. *Right:* The time-course on the right shows the mean (± SEM) peak OMR index scored by the three experimental groups before (pre) and after the injection. The OMR index of 1.2 is the reference value to discriminate between perceived (>1.2) and non-perceived (<1.2) pattern. The mean (± SEM) OMR index of vehicle-injected WT mice (green) over the two-weeks of observation is shown for comparison. **g.** Extracellular recordings of RGCs were performed with HD-MEA on retinal explants isolated from rd10 mice 2 days after the intravitreal injection of either DMSO or Ziapin2. Pie charts show the percentage of RGCs responsive to full-field cyan light stimulation (2 mW/mm^2^, 500 ms) in retinas from rd10 mice injected with either DMSO (top, black sector) or Ziapin2 (bottom, red sector), with respect to the total number of recorded RGCs. The absolute number of cells is reported in parenthesis. The box plots on the right show the number of responsive ON, ON-OFF, and OFF RGCs classified based on their response to light. Panels b-f: *p<0.05, ***p<0.001, one-way ANOVA/Dunnett’s tests *versus* “pre” within group; ^#^p<0.05, ^##^p<0.01, ^###^p<0.001, one-way ANOVA/Tukey’s tests *versus* rd10/Ziapin2; ^ΔΔ^p<0.01, ^ΔΔΔ^p<0.001, one-way ANOVA/Tukey’s tests *versus* WT/DMSO (WT/DMSO, n=7; rd10/DMSO, n=14; rd10/Ziapin2, n=18). Panel g: ****p<0.0001, Fisher’s exact test; ^##^p<0.01, ^###^p<0.001, two-way ANOVA/Holm-Šídák tests (n = 3 for both rd10 mice/DMSO and rd10 mice/Ziapin2). For exact p-values and source data, see the Source data file.

To investigate the effects of the intravitreal administration of Ziapin2 on pattern perception in blind rd10 mice, we evaluated the optomotor response (OMR) to moving grating patterns of varying spatial frequencies presented in a randomized sequence. Perception of moving patterns induces innate head movements in synchrony with the stimulus that are tracked and quantified as an OMR score that, above the 1.2 threshold, indicates normal visual performances ^61,62^. As expected, untreated WT mice scored an OMR index > 1.2 in the spatial frequency range between 0.05 and 0.3 cycles/degree, which was not affected by the intravitreal injection of vehicle (**Figure 7f, Figure S13c**). On the contrary, aged rd10 mice exhibited poor OMR responses (scores < 1) that were not affected by the injected vehicle. Strikingly, the visual performance of age-matched rd10 mice was significantly increased after intravitreal injection with Ziapin2, with OMR scores significantly higher than 1.2 that lasted for the whole 2-week period of observation (**Figure 7f**), as also shown by the individual performances of the animals in the three experimental groups (**Figure S13c**).

To verify that the *in vivo* visual restoration in blind mice was dependent on the effects of Ziapin2 observed *ex vivo*, we performed HD-MEA recordings on retinas obtained from rd10 mice treated with either vehicle or Ziapin2 two days after the intravitreal injection. In agreement with the visual rescue observed behaviorally, full-field cyan light stimuli were able to trigger AP firing in 16.6% of the recorded RGCs from *in vivo* Ziapin2-treated rd10 mice *versus* 1.7% of the recorded RGCs from vehicle-treated rd10 mice (Fisher’s exact test, p<0.0001; **Figure 7e**). Statistical analysis of the firing responses of RGC subclasses revealed a highly significant increase of responsive ON and ON-OFF RGCs in Ziapin2-treated as compared with vehicle-treated rd10 mice, with a smaller and non-significant increase in responsive OFF RGCs (**Figure S14g**).

In parallel with HD-MEA recordings, transversal sections of the retinas were subjected to immunohistochemistry to evaluate the depletion of rods and cones, as well as the possible presence of proinflammatory effects of Ziapin2 injection. No Ziapin2 fluorescence could be reliably detected because of the intrinsically weak fluorescence of the compound due to the energy photoconversion into the isomerization process ^50^ and its substantial dilution while reaching the retinal layers after intravitreal injection. The quantitative morphological analysis of cone-arrestin (**Figure S14a**) and rhodopsin (**Figure S14b**) revealed the total disappearance of rods and very scarce residual cone cell bodies devoid of external segments in the outer retina (**Figure S14d,e**), consistent with the lack of light responses. Notwithstanding Ziapin2 was previously demonstrated to be highly biocompatible and not to affect neuronal viability *in vitro* and *in vivo* ^48^, we analyzed GFAP and Iba1 immunoreactivities as markers of Muller cells/astrocytes and microglia activation, respectively, after intravitreal injection of either vehicle or Ziapin2 in WT and rd10 mice (**Figure S14c**). Consistent with previous results, no Ziapin2-induced increases in GFAP or Iba1 immunoreactivity/microglial cell number were observed in dystrophic rd10 mice, testifying to the absence of overt proinflammatory activity of Ziapin2 administration (**Figure S14f,g**).

## DISCUSSION

The use of membrane-targeted photochromic molecules to treat visual impairments is a promising strategic alternative to existing retinal prosthetics ^49,63^. We leveraged the interesting *Janus* properties of Ziapin2, a membrane-targeted photochromic compound that makes neurons hypoexcitable in the dark and phasically more excitable upon light stimulation by purely acting on the passive properties of the membrane, without interfering with ion channels or neurotransmitter receptors. The application of this compound to light-insensitive retinas affected by photoreceptor degeneration was inspired by the bifunctional behavior of the molecule that seemed well suited to mimic the ON/OFF processing mechanisms of the neuroretina. Currently, several materials and artificial retina devices can replace the phototransduction process of degenerated photoreceptors ^18,64,65^. However, the main challenge in using these materials and devices is the difficulty in recreating the natural encoding diversity found in normal retinas and indiscriminately imposing ON (light) signals on both ON and OFF cone BCs.

In this paper, we have found that Ziapin2 acts on retinal neurons with the same molecular mechanism described in artificial membranes, cell lines, and primary neurons, i.e., by modifying membrane capacitance in opposite directions depending on the dark/light condition. When directly puffed onto RGCs of WT retinas, Ziapin2 enhances RGC excitability in response to light, inducing RGC firing with an additive effect with respect to the physiological phototransduction mechanism. However, in fully degenerate retinas, in which the synaptic inputs to RGCs are insensitive to light, the individual RGC stimulation is insufficient to reach the firing threshold, and an action of Ziapin2 on second-order retinal neurons is necessary. When the whole retina was incubated with Ziapin2, cyan light activated multiple distinct RGC subpopulations (ON-sustained, ON-transient, ON-OFF, OFF-suppressed, OFF-transient RGCs), paralleled by the rescue of excitatory and inhibitory synaptic inputs onto RGCs, thus providing evidence that the critical effect of Ziapin2 is on second-order retinal neurons. These responses were always coherent with the stratification of the RGC dendrites in the inner plexiform layer, and the data obtained with specific synaptic blockers localized the Ziapin2 action at BCs with differential activation of cone ON and OFF BCs ^6–8^. While Ziapin2 stimulation could recover several distinct functional responses from RGCs, it was not possible to identify all the numerous functional subtypes of RGC present in healthy retinas ^1^ likely because of the advanced stage of degeneration of the rd10 mice and RCS rats used in the study, in the presence of a marked rewiring of the inner retina.

The mechanism of action of Ziapin2 is well suited to reactivate retinal processing. Being a membrane-targeted photoswitch, it works at single-cell resolution. The tonic increase in capacitance in the dark that is phasically abolished under cyan light stimulation represents a high-speed mechanism acting on the passive properties of the membrane and making neurons either reluctant or prone to excitation based on the presence of light. In the latter case, the light-induced sudden decrease in membrane capacitance boosts the power of any ionic current to change the membrane potential. Consistent with the transient changes in capacitance, the Ziapin2 effect is active also at relatively high frequencies of light stimulation, being still detectable at 15 Hz, considered the gold standard for “motion” vision. An additional effect of Ziapin2 would also be that to dampen the intense spontaneous oscillations that characterize degenerate retinas ^66–68^, thus lowering the background noise and increasing the signal-to-noise ratio and, therefore, signal discrimination. The Ziapin2’s capacitive mechanism can be potentially very effective in BCs, which mostly work through subthreshold oscillations of the membrane potential without generating APs. Moreover, BCs are distributed radially along the incoming light, whereas the other retinal neurons, excluding photoreceptors, are organized orthogonally to the light axis. As a result, BCs absorb more light per unit area due to this structural difference.

At the retinal network level, the action of Ziapin2 on the excitatory/inhibitory inputs to RGCs and the emergence of the physiological RGC subpopulations imply a differential activation of ON- and OFF-BCs. Ziapin2 partitions into the plasma membrane with high affinity for cholesterol-enriched lipid rafts and rapidly induces a ∼10-15% decrease in membrane capacitance upon light stimulation with respect to the dark value ^49^. Rod/cone-ON BCs are known to have longer axons that reach the most internal part of the internal plexiform layer (IPL; layers 3-5), while cone-OFF BCs have much shorter axons contacting OFF-RGCs in the external layers 1,2 of the IPL ^6–8,69,70^. Since only about 50% of cell capacitance is contributed by the soma ^71^, it is possible that rod/cone-ON BCs are more sensitive to the effects of Ziapin2 because of their higher overall cell capacitance and amount of Ziapin2 captured in the membrane. The larger absolute drop in capacitance experienced by rod/cone-ON BCs would induce a faster voltage change and a faster activation of rod/cone-ON BCs. Even if the percent capacitance change may not greatly differ between rod/cone ON-BCs and OFF-BCs, the light-evoked decrease in the membrane time constant will impact on the speed of electrotonic decremental propagation along the BC axon as a function of the axonal length. This can justify a more marked effect on ON-BCs exhibiting longer axons, while OFF-BCs with shorter axons will not be markedly affected. Alternative mechanisms for the selectively for ON-BCs could consist in a different enrichment of lipid rafts, for which Ziapin2 has high affinity ^49^ or of mechanosensitive channels that are expressed in retina neurons ^72^. Indeed, a recent report demonstrates that Ziapin2, by causing membrane deformation, modulates TRAAK mechanosensitive K^+^ channels overexpressed in HEK293 cells ^73^.

One potential mechanism for the distinct activation of ON- and OFF-BCs and relative RGCs could involve the light-induced activation of rod BCs and AII amacrine cells connected to OFF-BCs through a glycinergic synapse. This pathway was previously addressed to restore the ON-OFF responses by optogenetics ^74,75^. The experiments with strychnine to antagonize the AII amacrine cell inhibition revealed that, in the case of Ziapin2, this mechanism accounts for only about one third of the OFF responses of RGCs, and that other Ziapin2-dependent inhibitory mechanisms may exist to achieve the ON/OFF signal segregation. These observations are important in view of potential clinical applications. In fact, an involvement of rod BCs and AII amacrine cells located in the peripheral retina can extend the potential field of view for treated patients, while the existence of other inhibitory links may contribute to the rescue of the physiological ON/OFF signaling at the macular level. Finally, we cannot rule out a differential expression of voltage-gated channels in rod/cone-ON and cone-OFF BCs, and/or a more depolarized resting membrane potential in rod/cone-ON BCs in the absence of photoreceptor input ^49^. The preferential light-induced modulation of ON-BCs would then recreate the temporal tuning of excitatory and inhibitory inputs on RGC that generates the specific activity pattern of the distinct subpopulations of RGCs. Notably, all different classes of RGCs activated by Ziapin2 in degenerate retinas correspond to the morphological nature of RGCs regarding dendrite stratification in the IPL.

*Retinitis pigmentosa* can result from numerous pathogenic mutations of different genes encoding for proteins expressed by either photoreceptors or RPE cells, with distinct progression of the degeneration, making gene therapy for the definitive correction of the gene lesion very challenging. We demonstrated the rescue of light-evoked retinal circuit activity in rd10 retinas bearing a mutation in the photoreceptor-expressed gene *Pde6B* ^52,53^. At 6 months of age, these mice display a total absence of rods and very scarce residual cone cell bodies devoid of external segments, which have been shown to be at least two orders of magnitude less responsive than WT cones ^76^, consistent with the lack of light responses also at high luminances. To demonstrate that Ziapin2 can reactivate degenerate retinas irrespective of the causative mutation triggering photoreceptor degeneration, we also tested Ziapin2 in degenerate RCS retinas bearing a mutation in the RPE-expressed *Mertk* gene^54^ that impairs photoreceptor maintenance and survival and causes a rod-cone degeneration in humans ^77^. To this aim we used 10-month-old pink-eyed RCS rats in which rod/cone degeneration is complete at 6 months of age ^38,78^. Notably, we fully replicated the results obtained in the rd10 degeneration down to the effects on RGC firing, activation of distinct RGC subpopulations, activation of synaptic conductances, and response to synaptic blockers, demonstrating that Ziapin2 can be active on any retina circuit that has lost photoreceptors and light-sensitivity.

Administered *in vivo* to fully blind rd10 mice, Ziapin2 induced a stable rescue of light-driven behaviors and spatial discrimination up to two weeks after a single intravitreal injection by reactivating a large cohort of RGCs. Moreover, we confirmed that Ziapin2 is non-toxic and does not trigger proinflammatory effects in the retina, consistent with what we have previously shown in primary neurons and cerebral cortex ^49^. To administer Ziapin2, we used the minimally invasive intravitreal administration, widely used in patients, demonstrating that it could easily permeate the inner limiting membrane to reach inner retinal neurons and exert its physiological effects.

An alternative strategy to target the sole rod/cone-ON BCs is optogenetics, using the specific *Grm6* promoter ^79,80^ (for review, see ^81^). However, genetic targeting of rod/cone-ON BCs is far from optimal, particularly in degenerate retinas ^82^. As a result, no significant differences in visual restoration have emerged between genetic targeting of rod/cone-ON BCs and generalized opsin expression in RGCs independently of their phenotype ^65,83^. As a photochromic molecule, Ziapin2 can be benchmarked with other photo-pharmacological tools for neuron stimulation. Those pioneered by the Isacoff’s group rendered glutamate receptors photoswitchable by UV light but required a genetic modification of the channel to allow the covalent bond with the azobenzene-glutamate activator (for review, see ^84^). More recently, a whole battery of derivatized azobenzenes (including AAQ, DENAQ, BENAQ, DAD) pioneered by Kramer and collaborators and currently in Phase I/II clinical trials ^85^ were found to enter RGCs and mediate their light-induced activation by blocking K^+^ channels from the intracellular side (for review, see ^86^). While most of these molecules target the final retinal output irrespective of the physiological ON/OFF processing, DAD was reported to modulate, at least in part, the activity of BCs, by mostly eliciting ON responses and very rare OFF responses ^87^. On the contrary, Ziapin2 does not affect endogenous ion channels, primarily targets BCs, and reactivates excitatory and inhibitory synaptic currents in RGCs, promoting the re-emergence of physiological RGC subpopulations that can be essential to induce a sustained restoration of functional vision ^83^. In view of its potential therapeutic use, some limitations of Ziapin2 treatment remain, such as the lack of spectral sensitivity in the green/red region of the spectrum that may require the use of goggles, the partial solubility in physiological solutions, and the time limitations of the physiological effects due to membrane turnover ^49^. Further studies are ongoing to generate water soluble and red-shifted Ziapin2 derivatives and encapsulate the molecule for safe and prolonged delivery.

In conclusion, we have shown that Ziapin2, with its amphipathic nature, can intercalate into all retinal neurons and restore the physiologically complex network processing of visual information in degenerate retinas from preclinical models of RP. Not only did Ziapin2 restore the light sensitivity of fully degenerate retinas with different pathomechanisms, but it also reactivated the physiological ON/OFF processing responsible for contrast sensitivity and spatial discrimination, allowing the emergence of a variety of RGC responses resembling the normal retina. The striking ability of Ziapin2 to restore the selective activation of RGC subpopulations reveals that the molecule, characterized by an innovative photo-transduction mechanism, has the potential to outperform the existing prosthetic, optogenetic or photo-pharmacological treatments that are currently in clinical testing for RP therapy.

## Acknowledgements

The Authors thank Drs. M.M. La Vail (Beckman Vision Center, University of California San Francisco, CA) for kindly providing non-dystrophic RDY and dystrophic RCS rats; Luca Maragliano (IIT and Polytechnic University of Ancona, Italy) for providing the molecular dynamics snapshots shown in Fig. 1a; M. Cilli and L. Emionite (IRCCS Ospedale Policlinico San Martino, Genova, Italy) for assistance in the surgical procedures; R. Ciancio, I. Dall’Orto, A. Mehilli, R. Navone and D. Moruzzo (Istituto Italiano di Tecnologia, Genova, Italy) for technical assistance. The research was supported by Telethon-Italy (project # GMR22T2013), The Italian Ministry of Health (project Ricerca Finalizzata # GR-2021-12374630), H2020-MSCA-ITN 2019 “*Entrain Vision*” (project 861423), The Italian Ministry of University and Research (PRIN2020 project #2020XBFEMS) and IRCCS Ospedale Policlinico San Martino (Ricerca Corrente and 5x1000 grants).

## Author contributions

G.Z. and S.D.M. performed the *ex vivo* patch-clamp experiments; C.B. and V.S. designed and engineered Ziapin2; G.Z. and S.C. performed and analyzed the *in vivo* experiments. E.D. wrote the algorithms and analyzed the HD-MEA data. F.B. and S.D.M. conceived the work, planned the experiments, analyzed the data, and wrote the manuscript.

## Competing interests

The authors declare no competing interests. C.B., V.S., G.L., and F.B. are among the inventors of the patent application *"Photochromic compounds"* (PCT/IB2019/054530).

## Additional information

Supplementary information is available for this paper.

## Data availability

The experimental data supporting this paper’s figures and other findings are hosted at the Istituto Italiano di Tecnologia and can be accessed by contacting the corresponding authors. Source data are provided in this paper.

## Code availability

Custom-made code in Matlab and Igor Pro environments are hosted at the Istituto Italiano di Tecnologia and can be accessed by contacting the corresponding authors.

## Inclusion and Ethics

The Authors followed the recommendations of the “*Global Code of Conduct”* when designing, executing and reporting their research.

## MATERIALS AND METHODS

### Ethical approval and animal handling

All animal handling and experimental protocols complied with the guidelines established by the European Community (Directive 2014/26/EU of 4 March 2014) and were approved by the Italian Ministry of Health (Authorization # 357/2019-PR). Wild-type C57Bl6/J and homozygous rd10 mice on a C57Bl6/J background were used for this study. Breeders were obtained from The Jackson Laboratory (Bar Harbor, ME). Pink-eyed Royal College of Surgeons (RCS) dystrophic rats and non-dystrophic normal sighted controls (RDY) were kindly provided by Dr. M.M. La Vail (Beckman Vision Center, University of California San Francisco, CA). Animals were housed under standard conditions on a 12/12 light-dark cycle with *ad libitum* food and water. A balance between males and females of 6-month-old mice and 10-month-old rats was maintained with a random selection of the experimental groups.

### Preparation of retinal explants

Mouse retinal explants were obtained from 6-month-old WT and homozygous rd10 mice in the C57Bl6/J background. Rat retinal explants were obtained from 10-month-old RDY and RCS rats. Animals were dark-adapted for at least 60 min and euthanized by CO_2_ inhalation followed by cervical dislocation. The eyes were enucleated under dim red light, to preserve dark adaptation, and were transferred to a Petri dish containing carbo-oxygenated Ames’ medium (US Biological, Cat. N. A1372-25). After removal of the cornea, the eyecup was cut in half, and the retina was detached from the sclera. For patch-clamp recordings, the retina was placed photoreceptor side down in a recording chamber and transferred to an Eclipse FN1 microscope stage (Nikon Europe BV, Amstelveen, The Netherlands), illuminated with infrared light and viewed on a monitor connected with a CCD camera. The retina was continuously perfused with carbo-oxygenated Ames’ medium. To expose the RGC of interest, the above inner limiting membrane was removed with a patch pipette.

### Patch-clamp recordings and visual stimulation protocols

Whole-cell and cell-attached patch-clamp recordings were obtained from RGCs (soma diameter ≥ 20 µm) using an Axopatch 200B amplifier (Molecular Devices, San José, CA). For current-clamp recordings (I = 0) performed on normal sighted control and blind retinal explants, borosilicate patch pipettes (final resistance 4.5-6.0 MΩ) were filled with a solution containing (in mM): 126 K-gluconate, 4 NaCl, 0.02 CaCl_2_, 0.1 BAPTA, 0.1 EGTA-KOH, 1 MgSO_4_, 15 Glucose, 3 ATP-Na_2_, 0.1 GTP-Na, 5 HEPES, pH 7.3. To record light-evoked synaptic currents (blind retinal explants), cells were voltage-clamped at holding potentials ranging from -86 mV to +34 mV with voltage steps of 24 mV. For these recordings, patch pipettes (final resistance 5.0-7.0 MΩ) were filled with an intracellular solution containing (in mM): 120 Cs methane sulfonate, 4 NaCl, 0.02 CaCl_2,_ 1 MgSO_4_, 0.1 EGTA CsOH, 10 Phosphocreatine, 3 ATP-Na2, 0.1 GTP-Na, 10 HEPES, pH 7.3 with CsOH. Voltage-gated sodium currents were blocked by adding 5 mM QX-314 to the intracellular solution. The total light-evoked conductance was decomposed into excitatory and inhibitory components according to previously described methods ^88–91^.

For patch-clamp experiments performed on retinal explants isolated from normal sighted C57Bl6/J mice, basal light-evoked RGC firing activity was recorded in response to both cyan and green light flashes and compared to light-evoked responses to the same stimulation protocol after 2 min puff application of either 200 µM Ziapin2 or 10% DMSO in Ames’ medium by means of a second borosilicate glass pipette (final resistance 1.0-1.2 MΩ) positioned in the proximity of the patched RGC soma. For patch-clamp recordings of RGCs from blind retinal explants isolated from both homozygous rd10 mice and RCS rats, the retina was incubated for 30 min in the presence of 10 µM Ziapin2/0.5% DMSO immediately after the detachment from the sclera in 2-ml Eppendorf tubes with carbo-oxygenated Ames’ medium.

The Ziapin2- and light-induced changes in membrane capacitance (C_m_) and resistance (R_m_) of RGCs were measured by applying a voltage step of -5 mV, after the acute puff application of 200 µM Ziapin2. Pipette input resistance was always compensated. The capacitance current area (Q) was calculated using Origin software. C_m_ was calculated as C_m_ = Q/ΔV. Light-dependent changes in cell membrane capacitance were measured by illuminating with either green or cyan light (200 ms) during the 5-mV voltage step application and compared to the C_m_ obtained by the application of the same 5-mV voltage step in the dark. The same experimental protocol was used to measure membrane resistance (R_m_), calculated as R_m_ = ΔV/I.

Visual stimulation consisted of 500 ms flashes of chromatic circular spots (50 µm diameter) generated by a SPECTRA X Light Engine illumination source and focused through the microscope optics onto the patched RGC. The stimulus-evoked activity was recorded in response to different spectral outputs (cyan, ʎ = 470 and green, 544 nm) presented at subsequent increasing power densities (2, 10, 20, and 30 mW/mm^2^). Alexa Fluor 633 (50 µm) was added to both intracellular solutions for cell identification and morphology reconstruction.

### Analysis of electrophysiological recordings

Peristimulus time histograms (PSTHs) were built by recording light-evoked firing activity (50 ms bins: AP counts) in a time window starting from the onset of the light stimulus to 500 ms after light offset (ON cells) or, alternatively, from the offset of the light-stimulus to 1 s after (OFF cells). Light-evoked AP counts were averaged over 15 sweeps and normalized for the spontaneous AP counts (50 ms bins) in the 1.5 sec before light stimulation. The firing rate was computed using a kernel smoothing method with a Gaussian kernel (sigma = 25 ms) using a homemade routine with Python 3. Peak firing rate is defined as (A1-A2)/(A1+A2), where A1 is the maximal response amplitude measured in the dark and A2 is the maximal response amplitude measured either during the 500 ms light-stimulation (ON cells) or in a time-window of 1 sec after the end of light-stimulation (OFF cells). For AP waveform analysis, both the first spontaneous and the first light-evoked AP, after administration of Ziapin2/DMSO, were used to construct the plot (phase-plane plot) of the time derivative of voltage (dV/dt) *vs* voltage and were compared to the plot obtained from the first spontaneous AP under basal conditions (i.e. before Ziapin2/DMSO administration). The plot was used to extract the maximum rising slope, the maximum repolarizing slope, and the AP peak. For the V_m_ modulation analysis, whole-cell current clamp traces were analyzed by applying a lowpass filter in a time window of 1.5 sec starting from the onset of the 500-ms light stimulation; for each of the 15 sweeps, filtered mean V_m_ values were calculated over bins of 50 ms and normalized for the minimum (black squares, min) and the maximum (white squares, max).

### Morphological reconstruction of RGCs

At the end of patch-clamp experiments, retinas were fixed in 4% paraformaldehyde in phosphate-buffered saline (PBS) for 30 min at room temperature (RT) and then washed in 0.1 M PBS three times for 10 min each. Fixed retinas were then incubated in the presence of bisbenzimide (Hoechst 33342, 1:300; Sigma-Aldrich) for 30 min at RT and washed again (3 times, 10 min each), as previously described ^55^. Tissue samples were mounted with Mowiol (Sigma-Aldrich) between two glass coverslips. Whole-mount retinas were imaged using an SP8/HyD confocal microscope with super-resolution LAS-X Lightning deconvolution software (Leica Microsystems, Wetzlar, Germany).

### HD-MEA recordings and visual stimulation protocols

Extracellular recordings of light-evoked APs of RGCs were performed using a CMOS-based HD-MEA simultaneously recording from 4,096 electrodes (BioCam X, 3Brain, Wädenswil, Switzerland). For *ex vivo* experiments, retinas were incubated with 10 µM Ziapin2 in Ames’ medium before recording; for retinas treated by *in vivo* intraocular injection of Ziapin2, tissues were explanted two days after surgery and used directly for extracellular recordings. Freshly isolated retinas (see *Retinal Explant Preparation*) were cut in half, and the hemi-retinas were placed photoreceptor-side up onto the surface of the HD-MEA (Arena Chip, 3Brain) pre-coated with poly-DL-ornithine hydrobromide (5 mg/ml; Sigma-Aldrich) to improve the contact of RGCs with the recording electrodes. All the experiments were carried out in a dark room at RT. Light stimulation consisted of a full-field flash of either cyan or green light (250 and 500 ms) generated by a Lumencor Spectra-X Light Engine (Optoprim, Vimercate, Italy) illumination source flashed at 0.25 Hz. The light spot was focused on the tissue sample through a Z16 APO microscope (Leica Microsystems) to obtain a final power density ranging from 0.003 to 2 mW/mm^2^. RGC light-evoked responses were recorded over the repetition of 20-60 sweeps under the same experimental conditions (i.e., stimulus duration and power density). Acquired data were processed for spike sorting and peak detection by a Henning sorting-based algorithm ^92^ provided by a 3Brain routine. Incorrect sampling of noise signal was discarded by subsequent manual supervision of the elaborated data. Neurons increasing spiking activity in response to light stimulation above a firing threshold defined as “mean spontaneous firing rate plus 2 x SD” were classified as “*responsive RGCs*”; the remaining neurons were considered “*non-responsive*” and excluded from the analysis. ON, ON-OFF and OFF responses were calculated as increases in firing activity within the duration of the light stimulus or within 500 ms after its end, respectively and polarity was confirmed by manual supervision. To block synaptic inputs to RGCs and investigate the origin of the light-evoked firing activity, 10 µM 6-cyano-7-nitroquinoxaline-2,3-dione (CNQX; Tocris,Bristol, UK), 20 µM L-(+)-2-amino-4-phosphonobutyric acid (L-AP4, Tocris) or 10 µM strychnine hydrochloride (Sigma-Aldrich) were applied by perfusion with oxygenated Ames’ medium for 15, 5 and 10 min, respectively. A custom Matlab (MathWorks, Natick, MA) code was developed for further offline analysis of the acquired data.

### Intraocular injections

Six-month-old blind rd10 mice and aged-matched WT mice of either sex were anesthetized via inhalation of 1.5% isofluorane delivered through a nose cone while the animal was lying under the surgical microscope. Before injections, pupils were dilated with 1% tropicamide eye drops (VISUfarma SpA, Roma, Italy) and locally anesthetized with 4% benoxinate hydrochloride (Alfa Intes, Casoria, Italy) eyedrops. A 10 µl NanoFil syringe (NANOFIL, World Precision Instruments, Sarasota, FL) was filled with either 800 µM Ziapin2 or vehicle (40% DMSO in saline) and connected to an intraocular injection kit (IO-KIT, World Precision Instruments) with a 34-gauge blunt needle. Before injection, better access to the intravitreal space was obtained by piercing the conjunctiva with a 30-gauge needle. A microinjector (UMP3T-1, World Precision Instruments) was used to intravitreally inject 1 µl of the same solution in each of the two eyes at a controlled flow rate of 200 nl/s under the stage of a surgical microscope (M651 Leica Microsystems), yielding a final vitreal concentration of 200 µM Ziapin2 in 10% DMSO. Immediately after injection, tobramycin and dexamethasone (0.3% + 0.1%, TobraDex®, Alcon, Milano, Italy) and 0.2% carbomer (Lacrigel®, Dompè, Milano, Italy) solutions were applied to the eyes to prevent surgery-related infections and corneal dehydration, respectively. After ocular surgery, animals were kept alone in the dark for 1 h to avoid stress and discomfort and let them recover from anesthesia.

### Light-Dark box test

After intraocular injection, mice were tested for their light sensitivity employing an apparatus consisting of two communicating chambers: a “lightbox” with transparent walls and a “dark box” with black walls to create a protected hiding space. The test is based on the innate behavior of mice to avoid brightly illuminated areas, and the preference towards the dark compartment is an index of physiological light perception. The apparatus was placed in a dark experimental room, and animals were dark-adapted for 1 h before the test. The test started by placing the animal in the center of the “light compartment” in the dark for 3 min before a controlled 5-lux intensity light was turned on above the box. Video recording during the two illumination conditions allowed us to monitor the time spent by the animal in the whole light compartment and its central area. To assess the Ziapin2 efficacy in restoring the individual sensitivity to light, animals were subjected to the test 15 days before injection to compare the pre- and post-intraocular surgery performances.

### Optomotor response (OMR)

To investigate spatial/pattern perception of blind and WT-injected mice, we studied the OMR to moving patterns of varying spatial frequency. The OMR apparatus (PhenoSys GmbH, Berlin, Germany) consists of a cubic box with a central elevated circular platform on which unrestrained mice are placed, surrounded by four LCD screens. Visual stimuli include shifting grating patterns with white and black bars of different spatial frequencies projected to the four screens to create a virtual cylinder. A mirror on the box floor creates the optical illusion of infinite depth. In the central position, a normal-sighted animal moves the head in sync with the visual stimulation. A camera is placed above the animal to automatically track body/head movements in response to shifting grating patterns, while software uses this information to determine visual thresholds (OMR scores ^61,62^). We designed a virtual stimulation protocol to evaluate rd10 mice performance after Ziapin2 injection that included randomly presented spatial frequencies of 0.05, 0.1, 0.15, 0.2, 0.25, 0.3, 0.35, 0.375, 0.4, 0.45, and 0.5 c/deg. Each spatial frequency was presented for 60 s at a speed of 12°/sec and was interspersed with a full field gray stimulus for 3 s. Vehicle-injected aged-matched rd10 and WT mice were used as controls. The score of 1.2 was taken as a cut-off to quantitatively identify blind and light-sensitive animals: animals performing below this OMR index were considered not to perceive the specific spatial frequency. The highest OMR index scored after intravitreal injection for each experimental and control animal was compared to the best performance obtained before injection.

### Retina histochemistry

Retinas from mice treated in vivo with the intraocular injection of either Ziapin2 or vehicle were explanted two days after surgery and fixed in 4% paraformaldehyde (Sigma-Aldrich) in 0.1 M PBS overnight, washed in 0.1 M PBS, cryoprotected by passing a 15–30% sucrose scale, embedded in OCT freezing medium (Tissue-Tek; Qiagen, Milano, Italy) and cryo-sectioned at 25 μm using an MC5050 cryostat (Histo-Line Laboratories). Transversal sections were mounted on gelatin-coated glass slides and stored at -20 °C before processing. Sections were first incubated with 10% normal goat serum (NGS, Sigma-Aldrich) at RT for 1 h to block non-specific antibody binding, then incubated overnight at 4 °C with the following primary antibodies: rabbit anti-rhodopsin (1:500; Abcam, Cat. No. ab221664), rabbit anti-cone arrestin (1:500; Merck-Millipore, Cat. No. AB15282), guinea pig anti-GFAP (1:500; Synaptic Systems, Cat. No. 173011) and rabbit anti-Iba1 (1:250; Wako, Cat. No. 01919741). Alexa Fluor 488- and 568-conjugated secondary antibodies (Alexa 488 goat anti-guinea pig and Alexa 568 goat anti-rabbit; ThermoFisher) were diluted 1:100 and incubated at room temperature for 1 h. Slices were rinsed three times in 0.1 M PBS to eliminate excess antibodies and mounted with Mowiol (Sigma-Aldrich). Retinal sections were imaged with an SP8/HyD confocal microscope (Leica Microsystems). Acquisition parameters were kept constant throughout the imaging sessions for comparison purposes. The quantitative analysis was performed using cell counts for cone arrestin, rhodopsin, and Iba1 and as integrated immunoreactivity for GFAP using the ImageJ software (NIH, Bethesda, MD).

### Statistical analysis

The sample size needed for the planned experiments (*n*) was predetermined using the G*power software for the ANOVA test by considering an effect size = 0.25-0.40 with (type-I error) = 0.05, and 1-ß (type-II error) = 0.9, based on similar experiments and preliminary data. Experimental data are expressed either as means ± SEM or as box plots (center line, median; square symbol, mean; box limits, 25^th^-75^th^ percentiles; whisker length, min/max data points), with *n* as the number of neurons (*ex vivo*) or number of individual animals (*in vivo*). The normal distribution of experimental data was assessed using either the D’Agostino-Pearson’s or the Shapiro-Wilk’s normality test. Two-tailed paired Student’s *t*-test or Wilcoxon’s signed-rank test was used to compare two sample groups, depending on sample pairing and normality distribution. One- or two-way ANOVA was used to compare more than two normally distributed sample groups, followed by Dunnett’s, Tukey’s or Holm-Šídák post-hoc multiple comparison test. The one-way Kruskal-Wallis’ ANOVA was used to compare more than two non-normally distributed sample groups, followed by Dunn’s multiple comparison test. To perform the statistical analysis of contingency tables, the Fisher’s exact test was used. The criterion for statistical significance was p*<*0.05. Statistical analysis was performed using OriginPro2020 SR1 and GraphPad Prism 8.3.0.

## Notes

### Competing Interest Statement

The authors have declared no competing interest.

