## Supplementary Figures 1-14 for "A membrane-targeted photoswitch restores physiological ON/OFF responses to light in the degenerate retina"

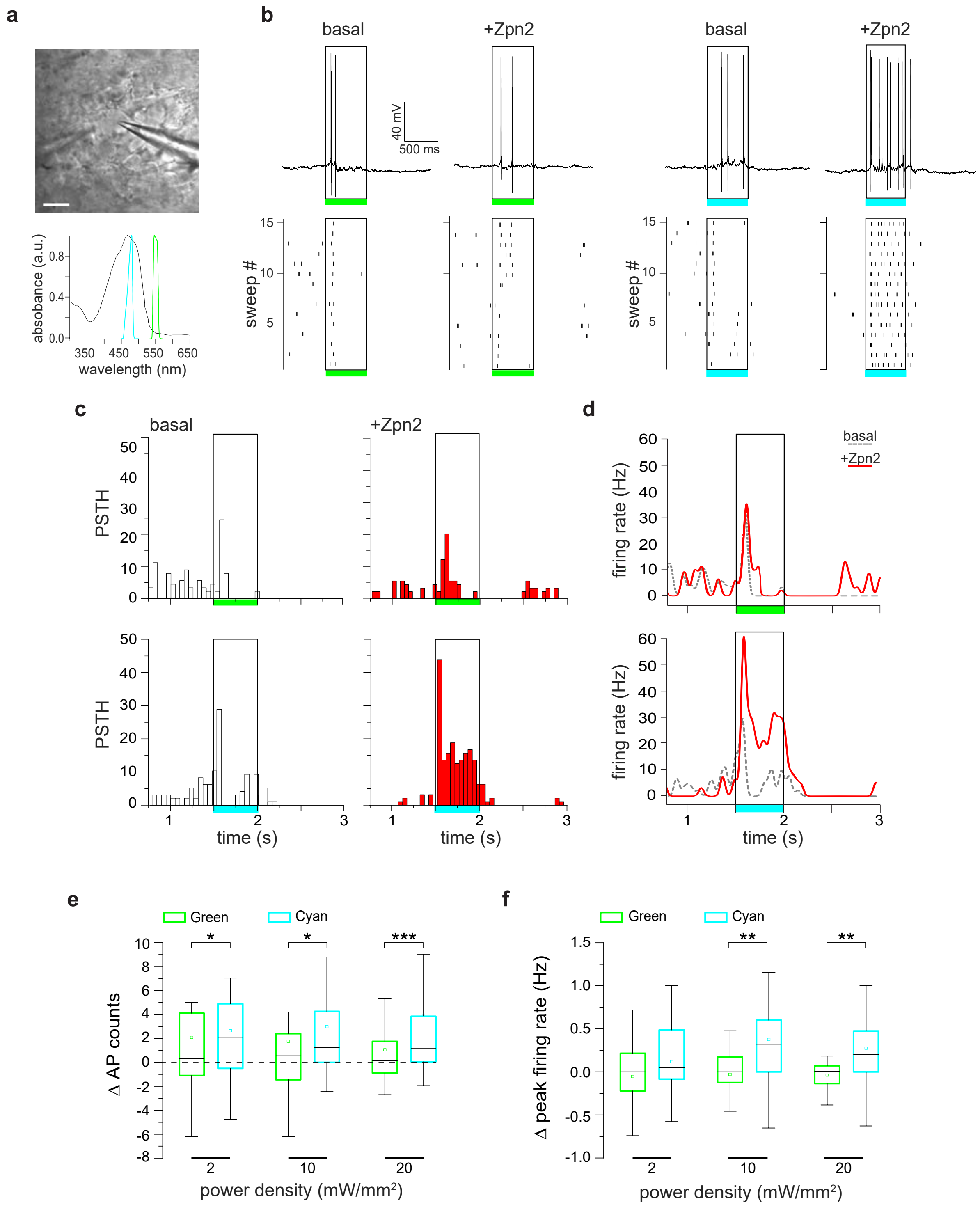


**Figure S1. The modulation of RGC firing by Ziapin2 in the WT retina is exclusively elicited by cyan light.**

**a.** *Top:* Representative bright-field image of a RGC showing a puff pipette (left) for the local application of Ziapin2 and a patch pipette (right) to record light-evoked firing activity. Scale bar, 20 µm. *Bottom:* Ziapin2 UV-Vis absorption spectrum (black trace) superimposed to the emission spectra of the cyan LED (ʎ = 470 nm) and the green LED (ʎ = 540 nm) laying outside the Ziapin2 spectrum.

**b.** *Top:* Representative current-clamp traces (I=0) recorded in the same RGC from a WT mouse retina in the absence (basal) or presence of Ziapin2 (Zpn2, 200 μM). Rectangles frame the responses during the light stimulus (500 ms, 20 mW/mm$^2$) applied with either green or cyan LED (colored horizontal bars). *Bottom:* Raster plots representing the firing activity of the same RGC as above over the repetition of 15 sweeps.

**c.** PSTHs of the same RGC, showing AP counts (50 ms-bins) recorded under basal conditions (white bars) or after the subsequent administration of Ziapin2 (red bars) and stimulated with either green or cyan light (open rectangle). An enhancement of firing activity was only evoked by cyan light stimulation.

**d.** Changes in the firing rate of the same RGC shown in **b** under basal conditions (dashed gray lines) and after the subsequent application of Ziapin2 (red line) in response to either green (top) or cyan (bottom) light-stimulation (open rectangle).

**e,f.** Box plots representing the light-evoked changes in AP counts (**e**) and peak firing rates (**f**) after acute Ziapin2 application with respect to the respective basal conditions. Box plots are shown for green and cyan light stimulation administered at increasing power densities (2, 10 and 20 mW/mm$^2$). Changes are expressed as the difference (Δ) between Ziapin2 puff application and the basal condition. *$p<0.05$; ***$p<0.001$; two-tailed paired Wilcoxon's signed-rank test green *versus* cyan light (n=34). For exact p-values and source data, see Source data file.

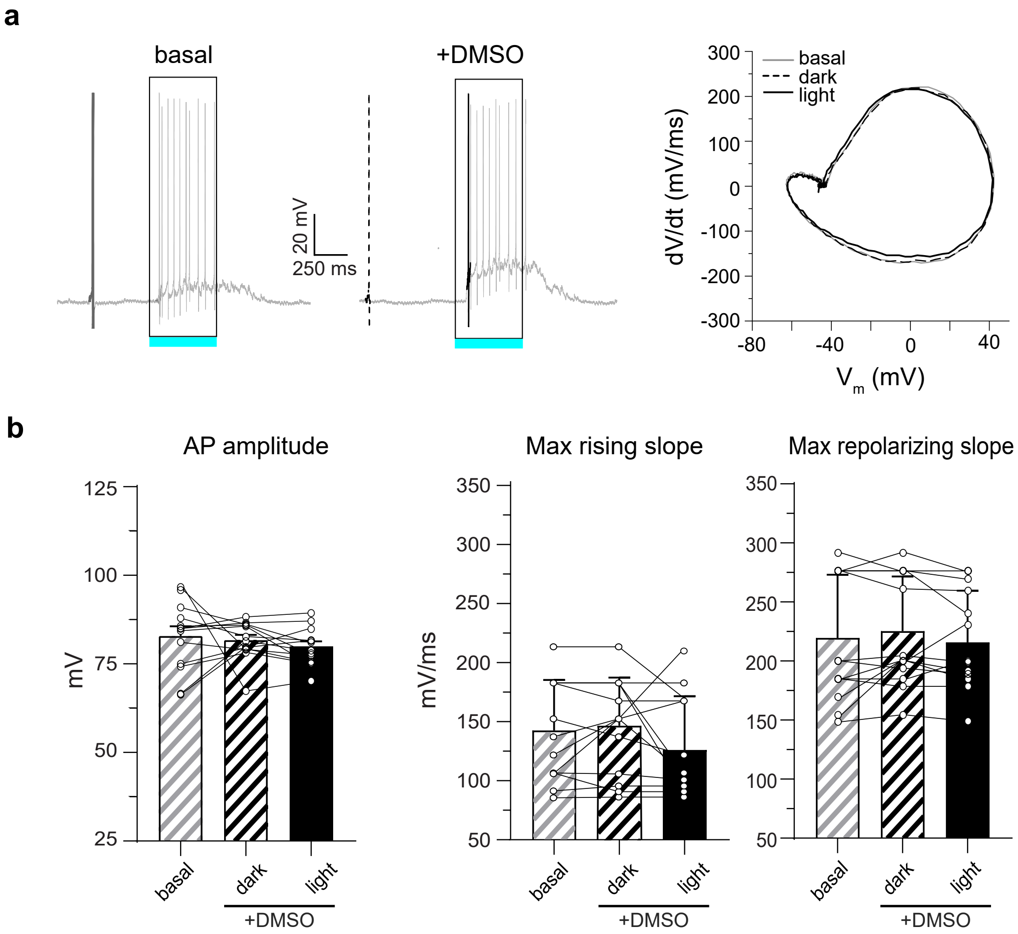


**Figure S2. DMSO does not affect AP amplitude and kinetics in RGCs from the WT retina.**

***a.*** *Left:* Representative whole-cell current clamp traces recorded in a RGC under basal conditions and after the acute application of DMSO (10% v/v). *Right:* The first spikes recorded under dark/basal conditions (gray line) and after DMSO either in the dark (dashed line) or in response to cyan light (solid line) were used for building phase-plane plots.

**b.** Changes in AP amplitude (left), maximal rising slope (middle) and maximal repolarizing slope (right) deduced from the phase-plane plot analysis of each recorded RGC before (basal) and after DMSO application in the dark and during subsequent cyan light stimulation (500 ms; 20 mW/mm$^2$). Bars represent means ± SEM with superimposed individual points. $p>0.05$; two-tailed paired Wilcoxon's signed-rank test (n=12). For exact p-values and source data, see Source data file.

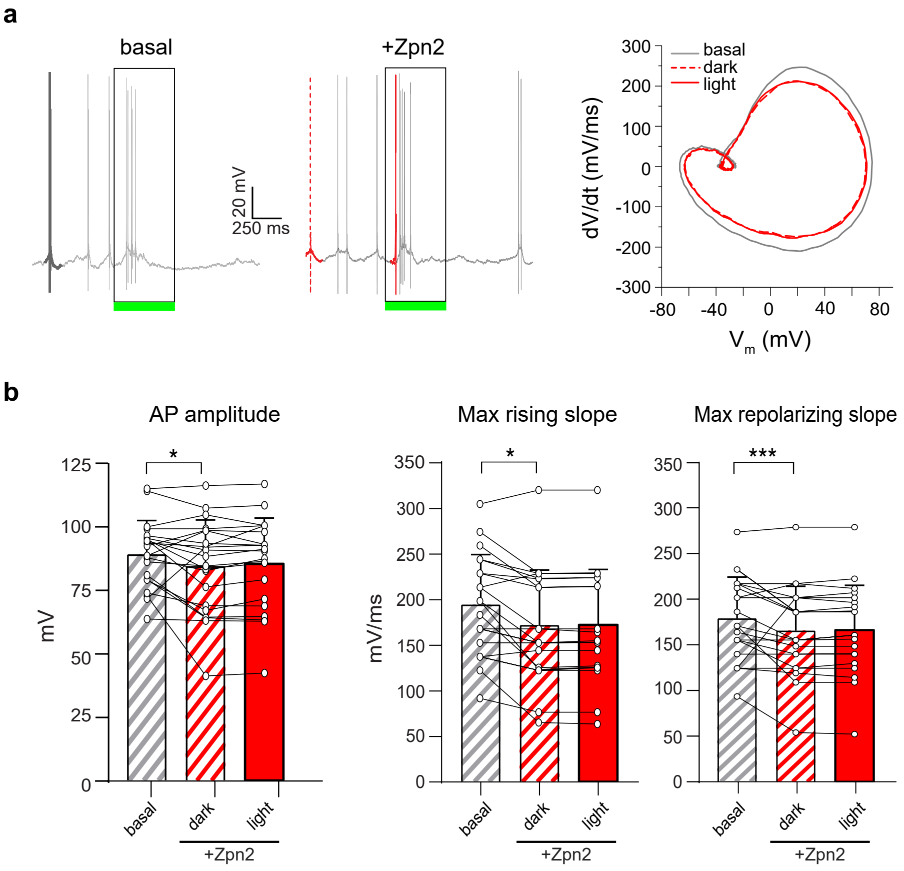


**Figure S3. Green light stimulation does not affect AP amplitude and kinetics in Ziapin2-treated RGCs from the WT retina.**

**a.** *Left:* Representative whole-cell current clamp traces recorded in the same RGC stimulated with green light under basal conditions (left) and after application of Ziapin2 (200 μM). *Right:* The first spikes recorded under basal conditions (gray line) and after acute application of Ziapin2 both in the dark (dashed red line) and in response to green light (solid red line) were used for building phase-plane plots.

**b.** Changes in AP amplitude (left), maximal rising slope (middle) and maximal repolarizing slope (right) deduced from the phase-plane plot analysis of each recorded RGC before (basal) and after Ziapin2 application in the dark and during green light stimulation (500 ms; 20 $mW/mm^2$). Bars represent means ± SEM with superimposed individual points. *$p<0.05$; ***$p<0.001$, two-tailed paired-sample Wilcoxon's signed ranked test (n=24). For exact p-values and source data, see Source data file.

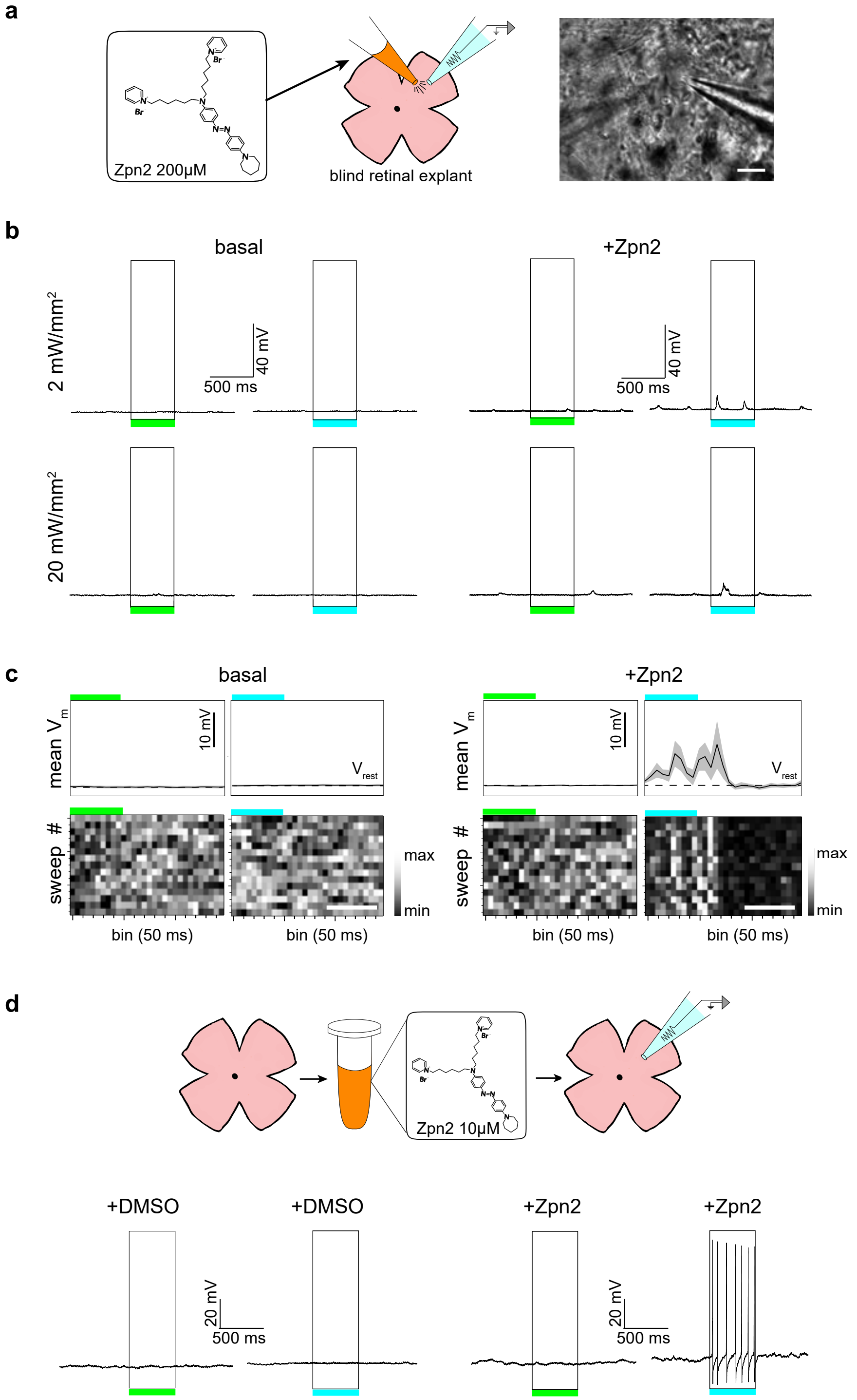


**Figure S4. Schematic of the experimental procedures for recording Ziapin2-treated blind retinal explants.**

**a.** Experimental design used to perform patch-clamp experiments on either WT or rd10 retinal explants. RGCs with a diameter of 15-20 μm are selected for recordings (right panel) and light-evoked responses are recorded upon either green or cyan illumination at 2 and 20 mW/mm$^2$ for 500 ms. After stimulation under basal conditions, a puff pipette is placed in the proximity of the patched RGC and used for the acute application of either vehicle (10% v/v DMSO) or Ziapin2 (200 μM) over 2 min before repeating the stimulation protocol with green/cyan light. Scale bar, 20 μM.

**b.** Representative whole-cell current-clamp recordings obtained in a RGC from blind rd10 retinal explant following the experimental design described in **a**. Under basal conditions (left), light stimulation with either green or cyan light did not induce either AP firing or membrane depolarization. Following application of Ziapin2 (right), significant cyan light-evoked membrane depolarizations, but not APs, were recorded. Light stimuli are represented by open rectangles labeled with green/cyan horizontal bars.

**c.** *Top*: Representative lowpass-filtered voltage traces recorded from the same RGC as in **b** under basal (left) and Ziapin2 (right) conditions in response to stimulation with either cyan or green light (horizontal bars). The plots display the mean (± SD, gray area) $V_m$ calculated within the same 1.5-s time window over 15 sweeps. *Bottom:* Grayscale representation of $V_m$ changes during the 15 sweeps (50-ms bins). Scale bars, 500 ms.

**d.** *Top:* Alternative experimental protocol to enhance Ziapin2 effects in blind rd10 retinas. Retina explants are incubated with either vehicle (0.5% v/v DMSO) or Ziapin2 (10 μM) for 30 min in carbo-oxygenated Ames' medium. After incubation, retinal explants are transferred to the stage of an upright microscope and perfused with Ames' medium for patch-clamp or HD-MEA recordings of RGCs. *Bottom:* Representative whole-cell current-clamp traces of a Ziapin2-treated RGC that responds with AP firing to cyan, but not to green, light stimulation. No effects of cyan/green light stimulation were observed in the DMSO-treated RGC. Light stimuli are represented by open rectangles labeled with green/cyan horizontal bars.

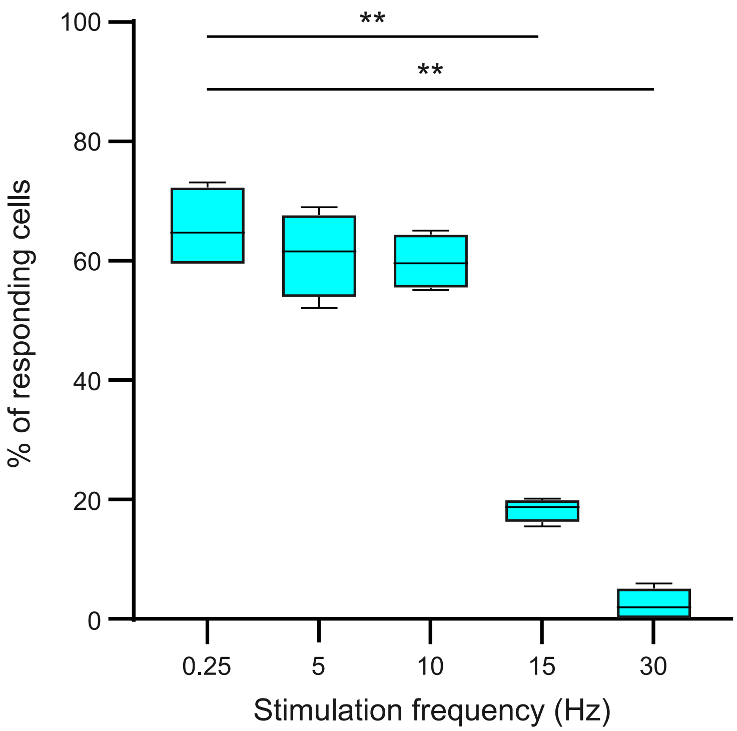


**Figure S5. RGC responses from blind rd10 retinas treated with Ziapin2 are evoked also by high-frequency cyan light stimuli.**

HD-MEA recordings were performed in rd10 blind retinal explants incubated with Ziapin2 (10 μM) and stimulated with pulsed cyan light (2 mW/mm$^2$) at 0.25, 5, 10, 15 and 30 Hz. The box plots show the percentage of responding RGCs as a function of the stimulation frequency. Cells were classified as responding when the FFT first harmonic was above the mean plus 2 x SD computed over the entire spectrum. **$p<0.01$, one-way ANOVA/Dunnett's tests (n = 4 rd10 mice). For exact p-values and source data, see Source data file.

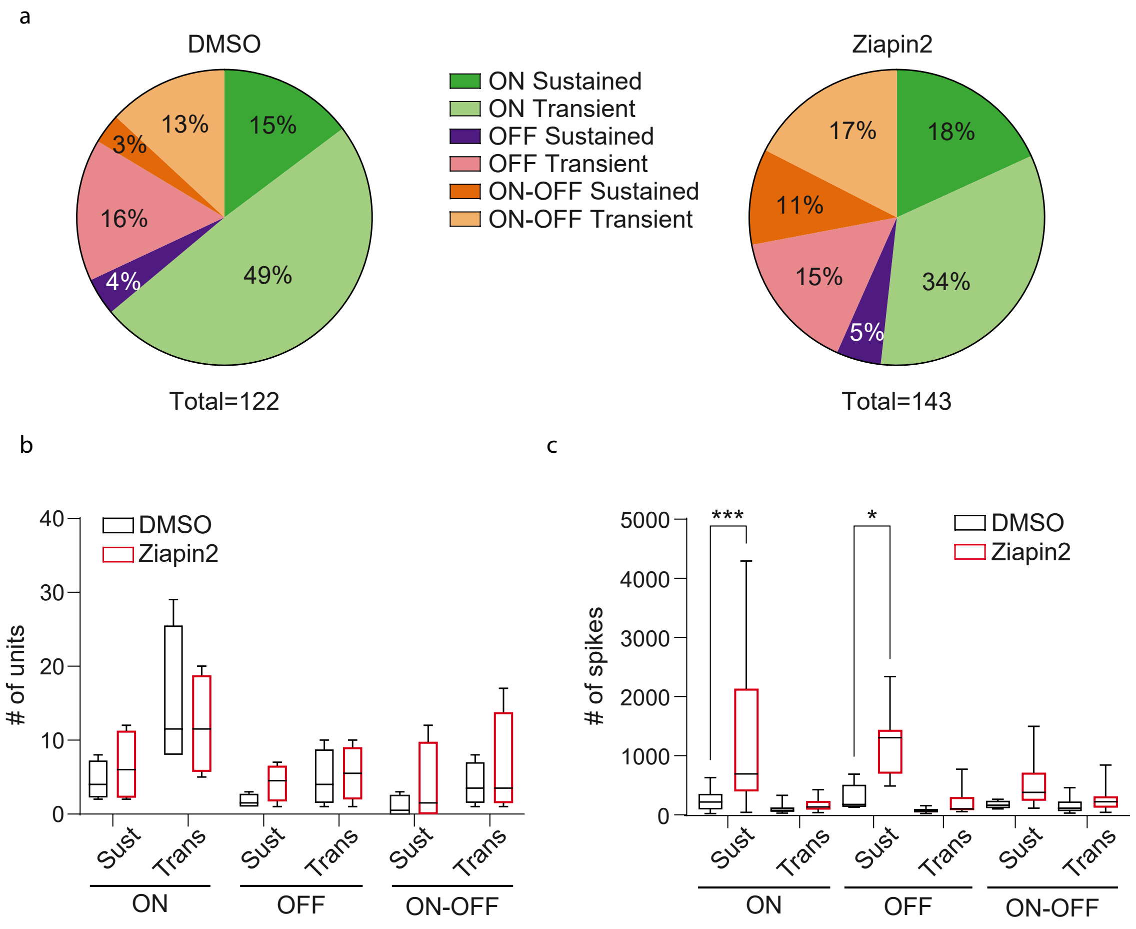


**Figure S6. Treatment of WT mouse retinas with Ziapin2 increases the firing rate of ON and OFF RGCs in response to cyan illumination.**

**a.** Pie charts of the overall percent distribution of sustained and transient ON, OFF and ON-OFF RGCs with respect to the total number of responding RGCs recorded by HD-MEAs in WT retinas incubated with either DMSO (0.5% v/v) or Ziapin2 (10 µM) and subjected to cyan light stimulation (250 ms, 2 mW/mm$^2$).

**b.** Box plots of the distribution of sustained and transient ON, OFF and ON-OFF RGCs in WT retinas incubated as described in **a** and subjected to cyan light stimulation. No significant changes in the distribution of RGC subclasses are observed.

**c.** Box plots showing the number of spikes generated by sustained and transient ON, OFF and ON-OFF RGCs in WT retinas incubated as described in **a** during cyan light stimulation. While Ziapin2 increases firing in all RGC classes, the effects are particularly clear-cut and significant on sustained ON and OFF RGCs. *$p<0.05$, ***$p<0.001$, one-way ANOVA/Tukey's tests (n = 4 WT mice for both DMSO- and Ziapin2-treated retinas). For exact p-values and source data, see Source data file.

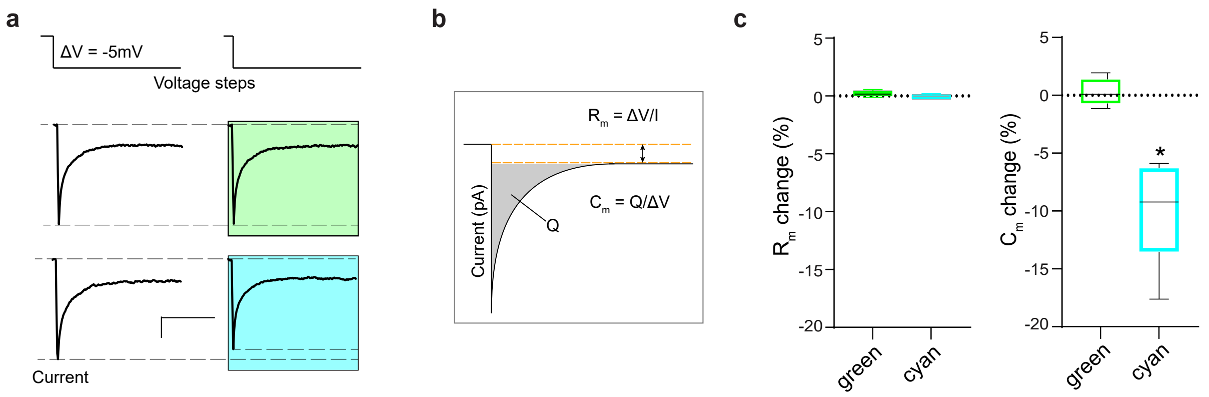


**Figure S7. Light-stimulation of Ziapin2-treated rd10 retinas decreases membrane capacitance of RGCs without affecting their membrane resistance.**

**a.** Representative current traces obtained in response to the application of -5 mV voltage steps to patched RGCs from rd10 retinas puffed with Ziapin2 (200 μM) before (*left*) and after (*right*) stimulation with either green or cyan light. Scale bars: 10 ms, 20 pA.

**b.** Schematic of the method used to extract $R_m$ and $C_m$ from the protocol shown in **a**. For further details, see Materials and Methods.

**c.** Box plots of the percent changes in $R_m$ and $C_m$ induced by either green (green boxes) or cyan (cyan boxes) light stimulation. *$p<0.05$, two-tailed paired Student's *t*-test ($n=10$). For exact p-values and source data, see Source data file.

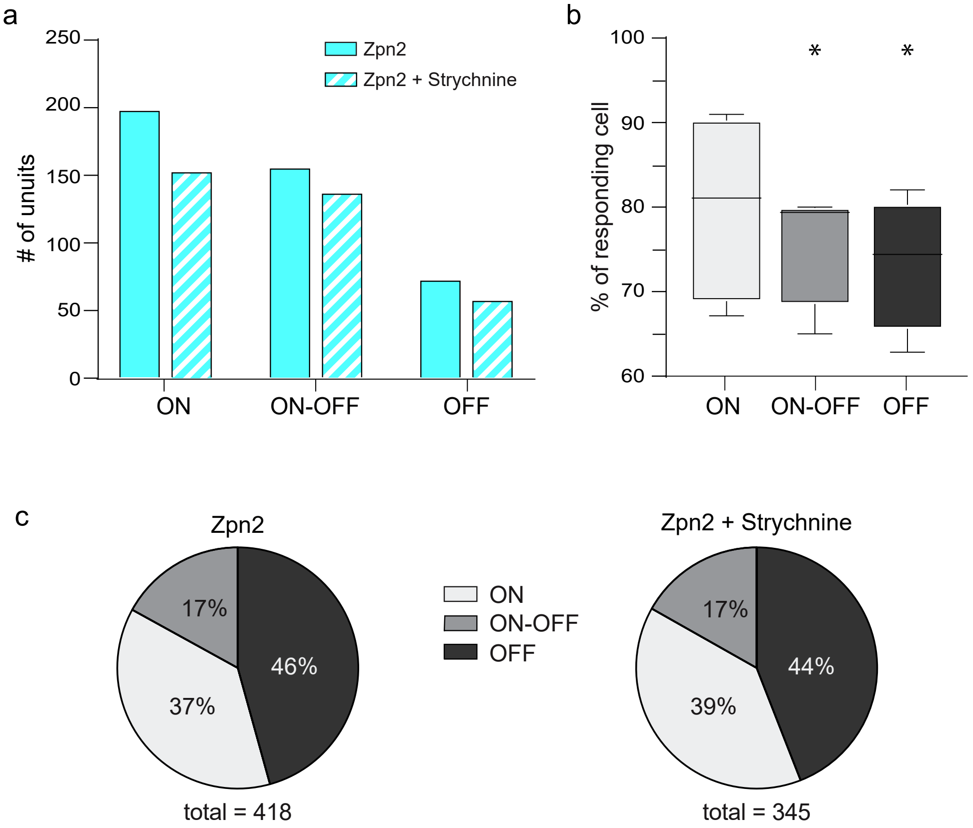


**Figure S8. Strychnine decreases OFF and ON-OFF RGC responses to cyan light in blind rd10 retinas incubated with Ziapin2.**

**a.** Explanted rd10 retinas incubated with Ziapin2 (10 μM) were recorded in HD-MEAs in the absence or presence of 10 μM strychnine to block glycinergic transmission between AII amacrine cells and cone OFF-BCs. The total number of ON, OFF and ON-OFF RGC units classified based on their response to light is shown.

**b.** The responding ON, OFF and ON-OFF RGCs after strychnine bath application are expressed in percentage of the respective number of responding RGCs under basal conditions.

**c.** Pie charts of the overall percentage distribution of ON, OFF and ON-OFF RGCs before and after strychnine bath application with respect to the total number of responding RGCs.

*$p<0.05$, one-way ANOVA/Dunnett's tests. For exact p-values and source data, see Source data file.

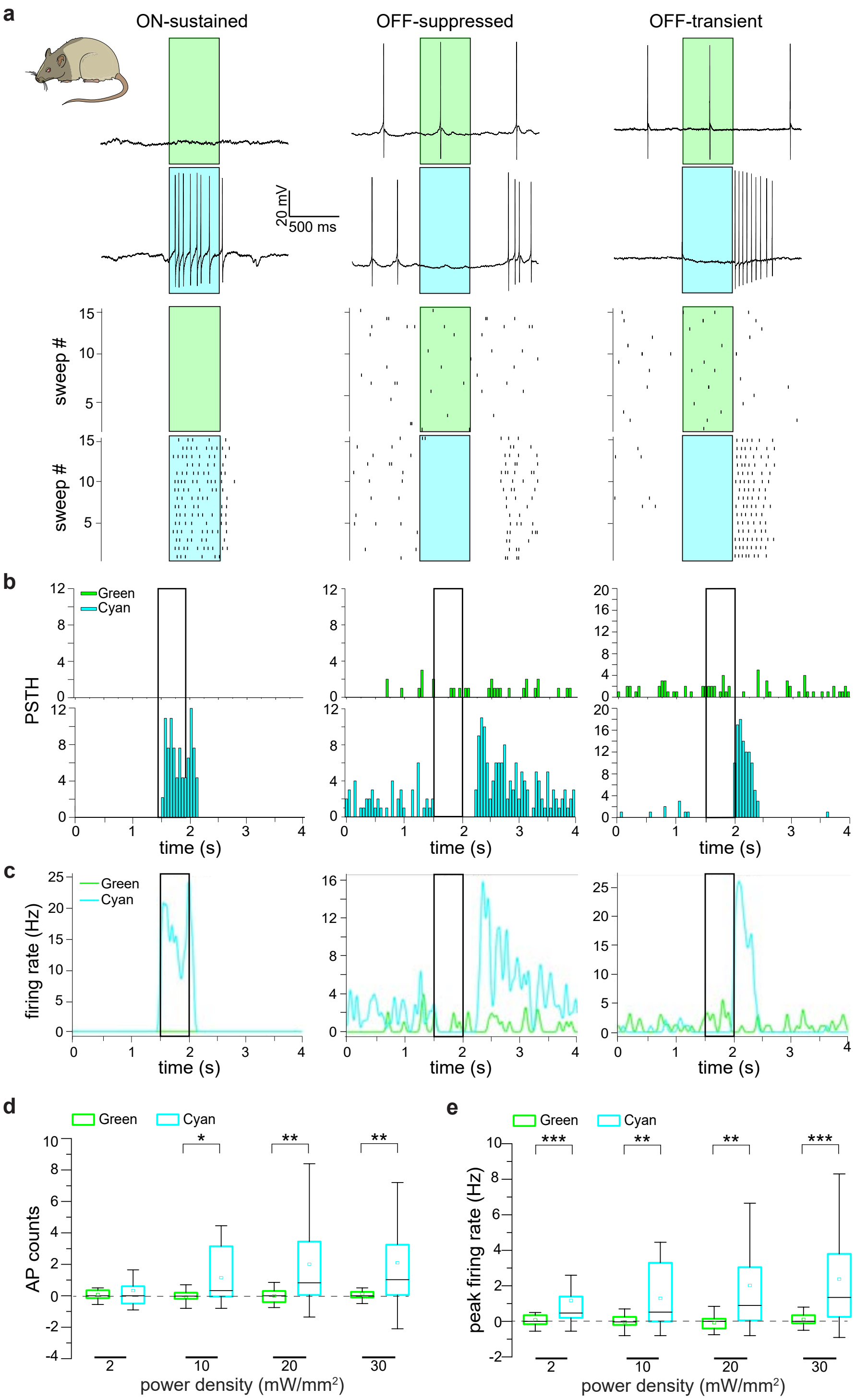


**Figure S9. Ziapin2 triggers light-dependent RGC firing in response to cyan light stimulation in advanced-stage RCS rats.**

**a.** *Top:* Representative responses of the RGCs in retinal explants from 10-month-old blind RCS rats to full-field illumination with either green or cyan light (500 ms; 20 mW/mm$^2$) after incubation in the presence of Ziapin2 (10 µM). Ziapin2 triggers light-evoked AP firing in response to cyan, but not green, stimuli

segregating the information into distinct subpopulations of RGC, originating ON- sustained, OFF-suppressed and OFF-transient responses. Light stimuli are shown as cyan- or green-shaded areas. *Bottom:* Raster plots of the voltage traces shown above over 15 consecutive sweeps.
**b,c.** Representative PSTHs (**b**) and instantaneous firing rate (**c**) of the RGCs recorded in **a**. In **b**, bars represent AP counts (50-ms bins) recorded in response to either green (top panels) or cyan (bottom panels) stimulation. Light stimuli (500 ms) are shown as open rectangles. No modulation of firing activity is evoked in response to green light. In **c**, the changes in the instantaneous firing rate in response to either green (green lines) or cyan (cyan line) stimulation are shown over time.
**d,e.** Box plots of the light-evoked AP counts (**d**) and peak firing rate (**e**) in all recorded RGCs in response to 500-ms stimulation with either green or cyan light (colored boxes) in a time window from the light onset to 1 s after light offset and normalized to the spontaneous activity recorded in the dark. Light stimulation was applied at increasing power densities (2, 10, 20, and 30 $mW/mm^2$) after 30-min incubation of blind retinal explants in the presence of Ziapin2 (10 μM). *$p<0.05$, **$p<0.01$; two-tailed paired-sample Wilcoxon's signed-rank test (n=22). For exact p-values and source data, see Source data file.

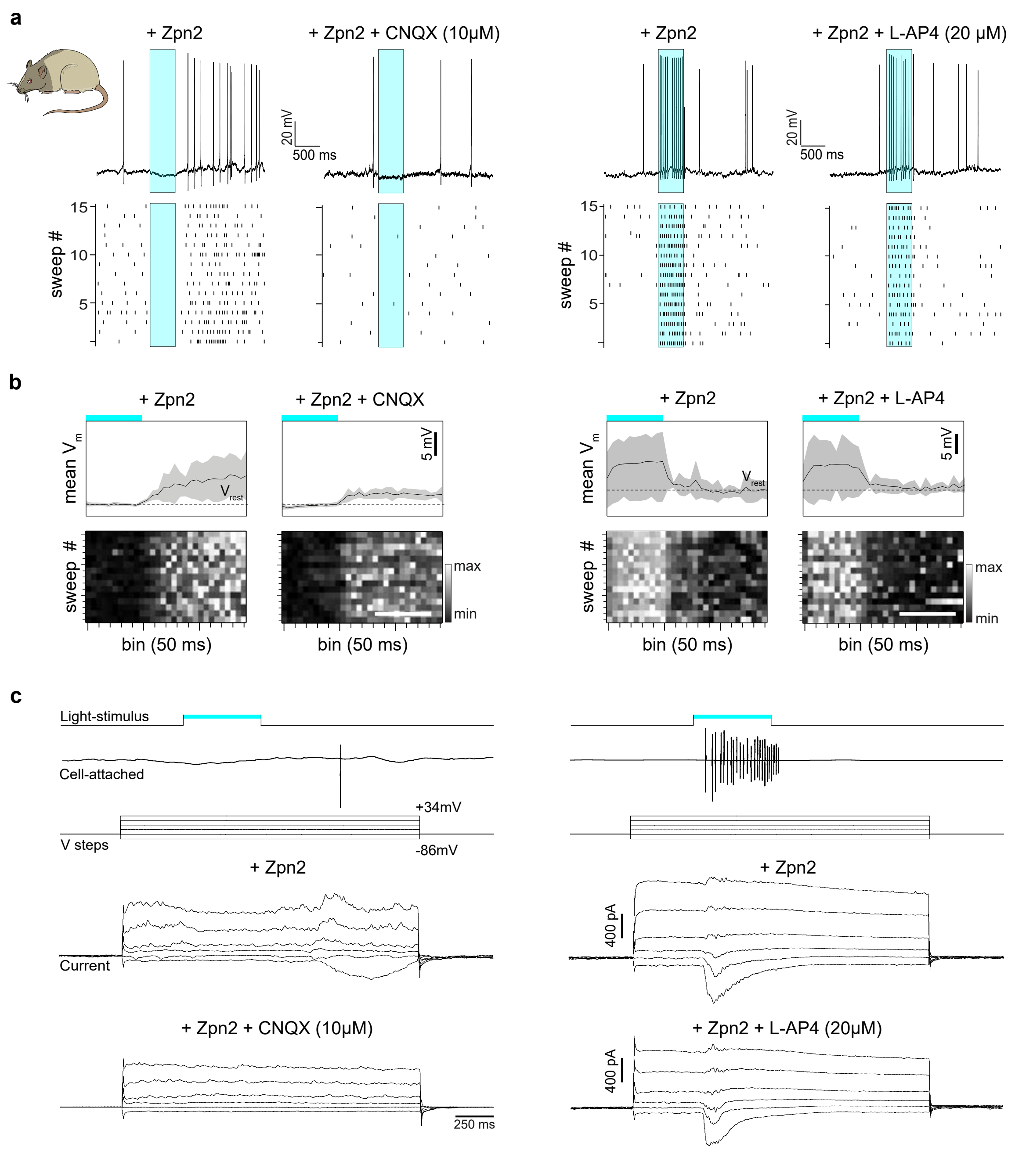


**Figure S10. Cyan light-evoked RGC responses in blind RCS retinas treated with Ziapin2 are generated in the inner retinal network.**

**a.** Representative whole-cell patch-clamp traces (top row) and relative raster plots (bottom row) of RGCs from retinas explanted from 10-month-old blind RCS rats incubated with Ziapin2 (10 µM) and interrogated with cyan light stimuli (20 mW/mm$^2$; 500 ms; cyan rectangles). *Left:* Light-evoked responses of an OFF-RGC before and after perfusion with the AMPA/kainate receptor antagonist CNQX (10 µM). *Right:* Light-evoked responses of an ON-RGC before and after perfusion with the mGluR6 receptor agonist L-AP4 (20 µM). L-AP4 did not alter light-induced firing activity, indicating that the response is not mediated by photoreceptor activation.

**b.** The same Ziapin2-treated RGCs as in **a** were stimulated with cyan light stimuli (20 mW/mm$^2$; 500 ms) before and after perfusion with either CNQX (*left*) or L-AP4 (*right*). *Top:* Representative whole-cell lowpass-filtered voltage traces in a time-window of 1.5 sec from the onset of the light stimulus. The plots display the mean (± SD, gray area) $V_m$ calculated within the same time-window over 15 sweeps. *Bottom:* Grayscale representation of $V_m$ changes during the 15 sweeps (50-ms bins).
**c.** *Top:* Representative voltage-clamp recordings in cell-attached configuration of a Ziapin2-treated OFF-RGC (*left*) and ON-RGC (*right*) subjected to full-field stimulation with cyan light (20 mW/mm$^2$; 500 ms; horizontal bars). *Bottom:* Light-induced currents recorded in the same RGCs after reaching the whole-cell configuration by applying voltage steps from -86 to 34 mV. On the left, light stimulation of the RGC elicited an OFF response, whose input currents are abolished after application of CNQX. On the right, the ON response of the RGC to cyan-light was generated by synaptic inputs that were not affected by L-AP4.

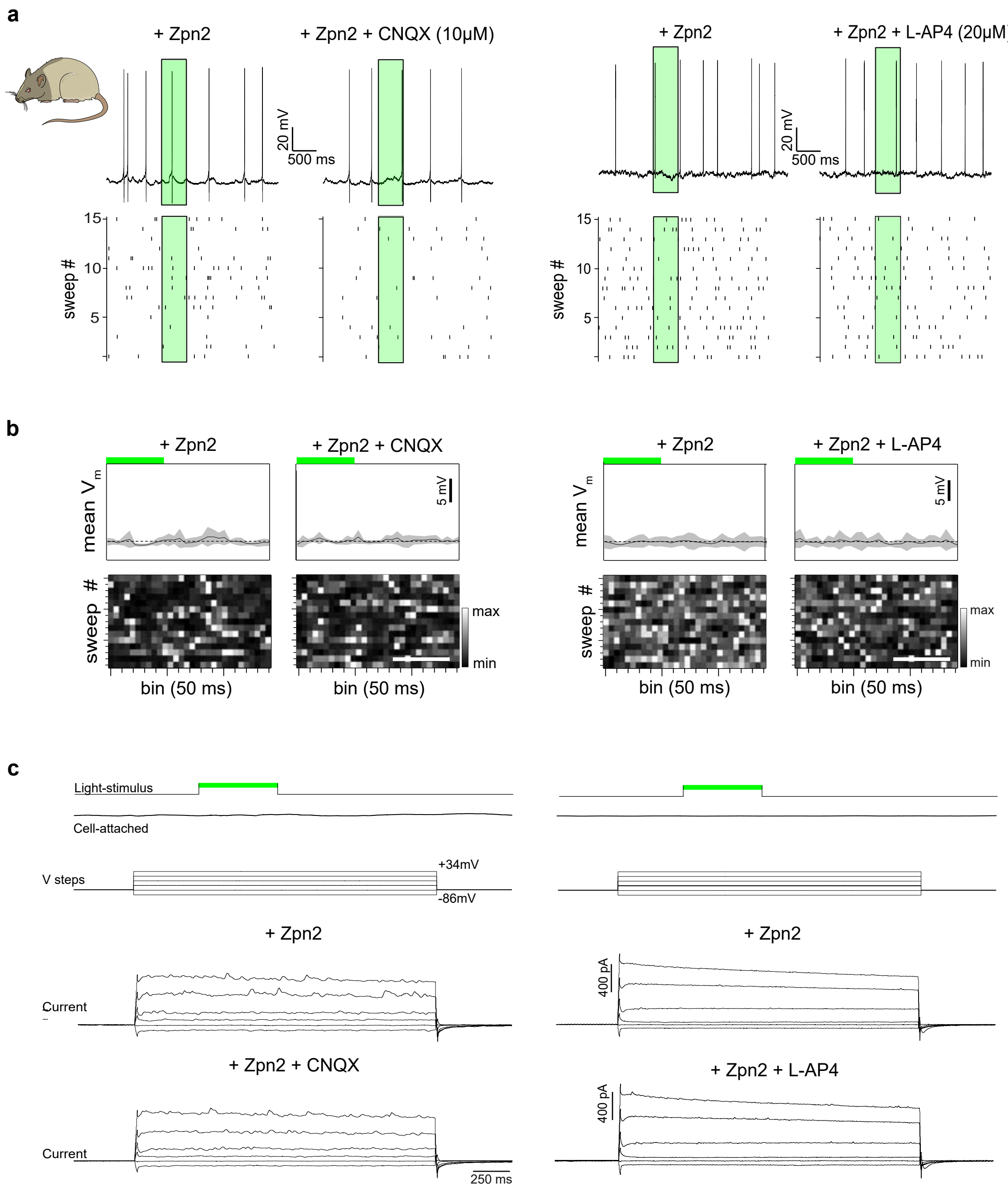


**Figure S11. Firing activity and synaptic conductances of Ziapin2-treated RCS RGCs are not altered by green light stimulation.**

**a.** Representative whole-cell patch-clamp traces (top row) and relative raster plots (bottom row) of RGCs from retinas explanted from 10-month-old blind RCS rats incubated with Ziapin2 (10 μM). *Left:* Responses to green light stimulation (20 mW/mm$^2$; 500 ms; green rectangle) before and after perfusion with CNQX (10 μM). *Right:* Responses of another RGC to green light stimulation (green rectangle) before (left trace) and after (right trace) perfusion with L-AP4 (20 μM).

**b.** *Top:* Lowpass-filtered voltage traces recorded from the same RGCs shown in **a** (top panels) in response to 500-ms green light stimulation (horizontal bars). Retinas were incubated for 30 min with Ziapin2 (10 μM) and recorded before and after perfusion with either CNQX (left panels) or L-AP4 (right panels). Panels display the mean (± SD, gray area) $V_m$ calculated within the same time-window over 15 sweeps. *Bottom:* Grayscale representation of $V_m$ changes in response to 500-ms green light stimulation during 15 sweeps (50-ms bins). Scale bars, 500 ms.

**c.** *Top:* Representative voltage-clamp recordings in cell-attached configuration of two RGCs from blind RCS retinal explants. After 30-min incubation with Ziapin2 (10 μM), green light stimulation (top trace) did not elicit any response. *Bottom:* Currents recordings performed in the same RGCs after reaching whole-cell configuration and applying voltage steps from -86 mV to 34 mV. Full-field stimulation with green light (20 $mW/mm^2$; 500 ms) was applied in the absence or presence of the synaptic blockers CNQX (10 μM; left traces) and L-AP4 (20 μM; right traces).

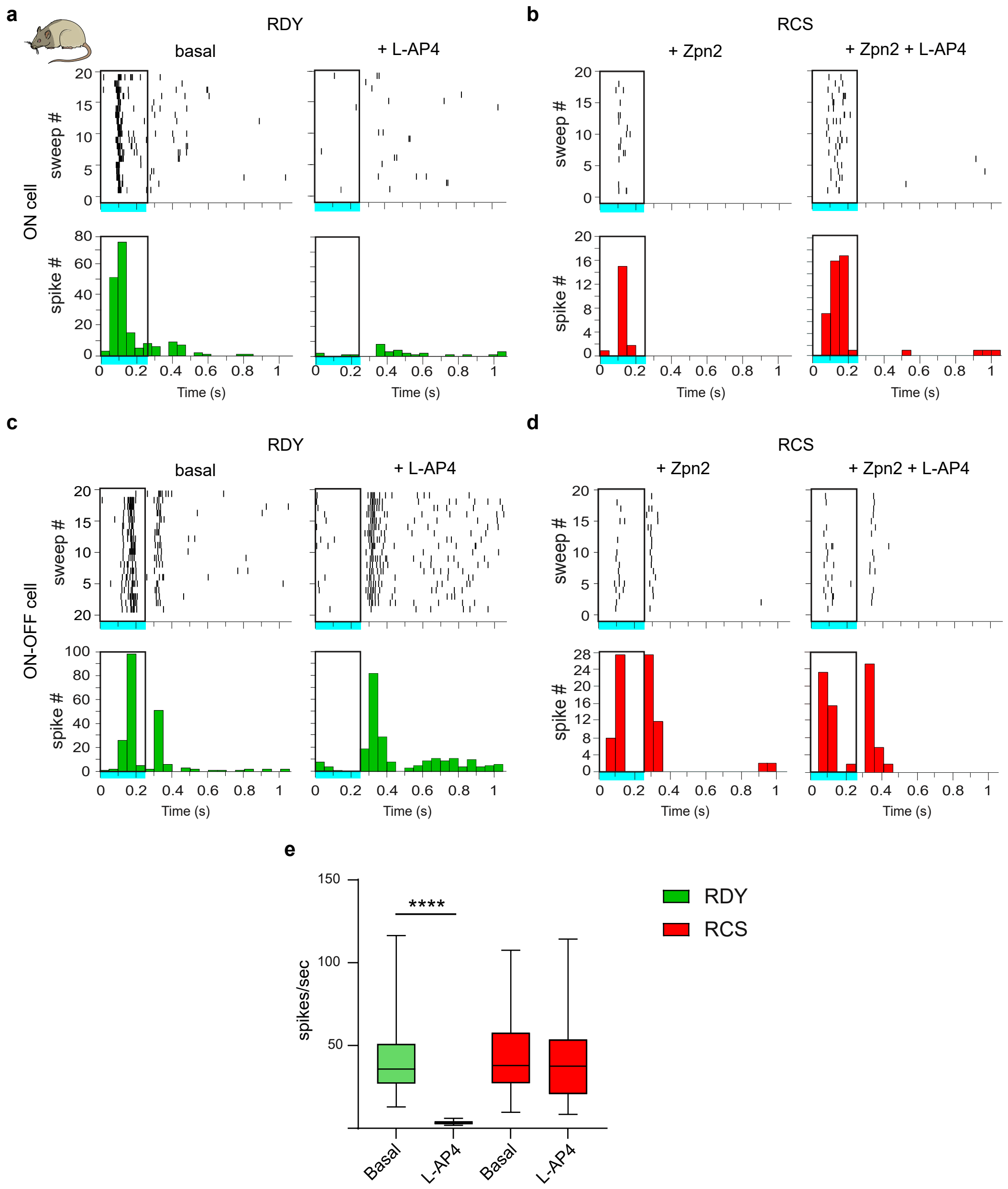


**Figure S12. Ziapin2 triggers ON-responses in blind RCS retinas by reactivating the inner retinal network.**

**a-d.** Raster plots (upper panels) and relative averaged PSTHs over 20 sweeps (lower panels) obtained from representative RGCs from non-dystrophic (**a**,**c**) and dystrophic (**b**,**d**) retinas explanted from 10-month-old RDY and RCS rats, respectively, and recorded with HD-MEAs. Untreated RDY retinas and Ziapin2-treated (10 μM) RCS retinas were stimulated with full-field cyan light (250 ms, 2 $mW/mm^2$; open rectangle) in the absence or presence of L-AP4 (20 μM). L-AP4 blocked the ON responses in RDY ON-RGCs, but not the OFF component in RDY ON-OFF RGC. The ON responses of Ziapin2-treated RCS RGCs are resistant to L-AP4, indicating that they result from the reactivation of the inner retinal network.

**e.** Box plots of the effects of L-AP4 on RGC firing in non-dystrophic RDY retinas and dystrophic RCS retinas (n=3 for both RDY and RCS rats). ****p<0.0001, two-way ANOVA/Tukey's tests. For exact p-values and source data, see Source data file.

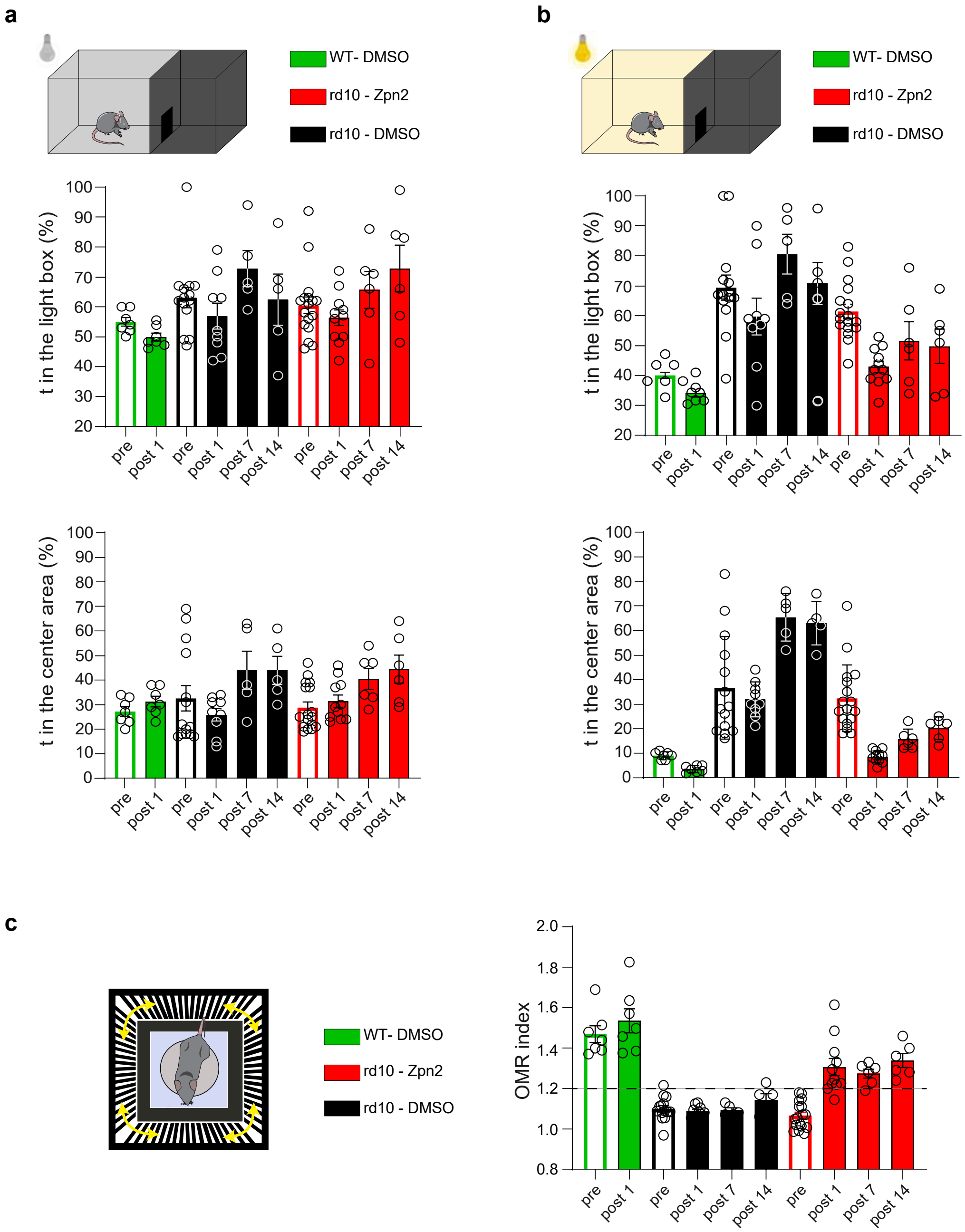


**Figure S13. Light-dark box and OMR behavior at various times after the intravitreal administration of either DMSO or Ziapin2.**

**a.** *Top:* Light-Dark box apparatus entirely kept in the dark. Animals were tested for 1 week before and during the 14 days after the ocular injection. *Middle:* Mean (± SEM and individual experimental points) percent time spent in the left compartment over the total duration of the test by WT mice (green) and age-matched blind rd10 mice treated with either vehicle (10% DMSO final vitreal concentration; black) or Ziapin2 (800 μM; red) in saline (1 μl/eye). Pre, before ocular surgery; Post, days after intravitreal injection. *Bottom:* Mean (± SEM) percent time spent in the center area of the left compartment.

**b.** *Top:* Light-Dark box apparatus during light stimulation (5 lux) in the left compartment. *Middle:* Mean (± SEM and individual experimental points) percent time spent in the light compartment over the total duration of the test by the three experimental groups. *Bottom:* mean (± SEM) percent time spent in the center area of the left compartment.

**c.** *Left:* Schematic representation of the Optomotor Response (OMR) apparatus. The unconstrained mouse instinctively follows the grating patterns rotating around it (yellow arrows) with synchronized head movements. *Right:* Bar plots show the means (± SEM and individual experimental points) of the peak OMR index scored by the three experimental groups before (pre) and over 2 weeks after the injection. The OMR index of 1.2 is the reference value to discriminate between perceived (>1.2) and non-perceived (<1.2) pattern. (WT/DMSO, n=7; rd10/DMSO, n=14; rd10/Ziapin2, n=18)

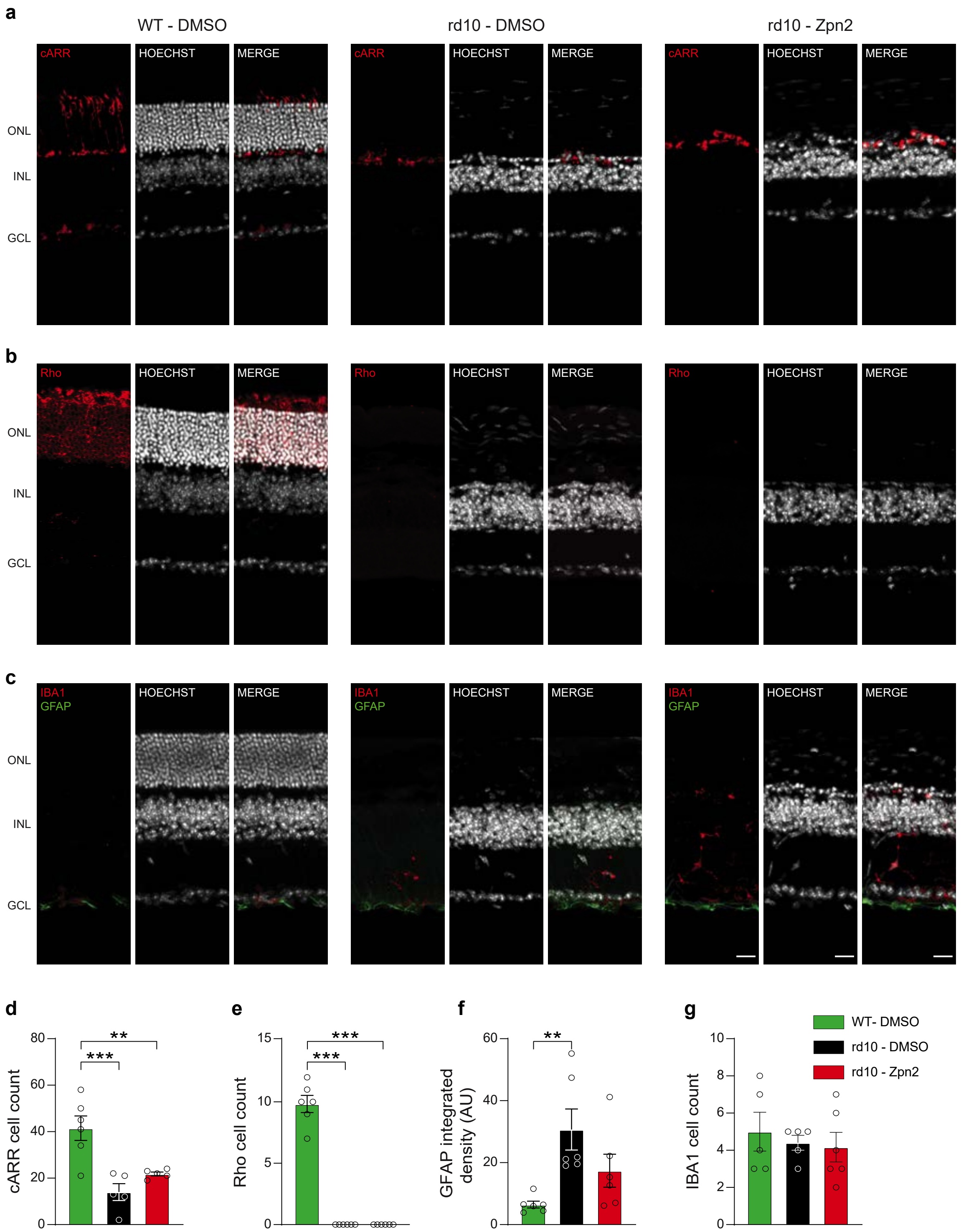


**Figure S14. Intravitreal Ziapin2 does not induce retinal inflammation.**
**a-c.** Transversal sections of the retinas were subjected to immunohistochemistry with antibodies to: (i) cone arrestin (cARR; **a**) and rhodopsin (Rho; **b**) to evaluate the depletion of cones and rods, respectively; (ii)

GFAP and Iba1 (**c**) to assess the potential proinflammatory effects due to Ziapin2 intravitreal injection on astrocytes and microglial cells, respectively.

**d,e.** Quantitative morphological analysis of cone arrestin (**d**) and rhodopsin (**e**) reveals the photoreceptor depletion in rd10 retinas.

**f,g.** Quantitative analysis of GFAP (integrated density; **f**) and Iba1 (cell counts; **g**) immunoreactivities shows the absence of proinflammatory effects of the intravitreal administration of Ziapin2 in dystrophic rd10 retinas. Kruskal-Wallis' ANOVA/Dunn's tests. **$p<0.01$; **$p<0.01$ ($n=5$-6 per experimental group).
